# Genomics of the “tumorigenes” clade of the family *Rhizobiaceae* and description of *Rhizobium rhododendri* sp. nov.

**DOI:** 10.1101/2023.01.02.522471

**Authors:** Nemanja Kuzmanović, George C. diCenzo, Boyke Bunk, Cathrin Spröer, Anja Frühling, Meina Neumann-Schaal, Jörg Overmann, Kornelia Smalla

**Affiliations:** Julius Kühn-Institut (JKI), Federal Research Centre for Cultivated Plants, Institute for Plant Protection in Horticulture and Urban Green, Braunschweig, Germany; Department of Biology, Queen’s University, Kingston, Ontario, Canada; Leibniz Institute DSMZ-German Collection of Microorganisms and Cell Cultures, Braunschweig, Germany; Microbiology, Technical University of Braunschweig, Germany; Julius Kühn-Institut (JKI), Federal Research Centre for Cultivated Plants, Institute for Epidemiology and Pathogen Diagnostics, Braunschweig, Germany

**Keywords:** crown gall, rhododendron, blackberry, taxonomy, genomics, pan-genome analysis

## Abstract

Tumorigenic members of the family *Rhizobiaceae*, known as agrobacteria, are responsible for crown and cane gall diseases of various agricultural crops worldwide. Tumorigenic agrobacteria are commonly found in the genera *Agrobacterium*, *Allorhizobium*, and *Rhizobium*. In this study, we analyzed a distinct “tumorigenes” clade of the genus *Rhizobium*, which includes the tumorigenic species *Rhizobium tumorigenes*, as well as strains causing crown gall disease on rhododendron. Here, high quality, closed genomes of representatives of the “tumorigenes” clade were generated, followed by comparative genomic and phylogenomic analyses. Additionally, phenotypic characteristics of representatives of the “tumorigenes” clade were analyzed. Our results showed that the tumorigenic strains isolated from rhododendron represent a novel species of the genus *Rhizobium* for which the name *Rhizobium rhododendri* sp. nov. is proposed. This species also includes additional strains originating from blueberry and Himalayan blackberry in USA, whose genome sequences were retrieved from GenBank. Both *R. tumorigenes* and *R. rhododendri* contain multipartite genomes, including a chromosome, putative chromids, and megaplasmids. Synteny and phylogenetic analyses indicated that a large putative chromid of *R. rhododendri* resulted from the cointegration of an ancestral megaplasmid and two putative chromids, following its divergence from *R. tumorigenes*. Moreover, gene clusters specific for both species of the “tumorigenes” clade were identified, and their biological functions and roles in ecological diversification of *R. rhododendri* and *R. tumorigenes* were predicted and discussed.

## 1 | INTRODUCTION

The family *Rhizobiaceae* contains genetically and phenotypically diverse bacteria isolated from various environments. Accordingly, *Rhizobiaceae* members exhibit remarkably diverse lifestyles, ranging from plant symbionts (rhizobia) and pathogens (agrobacteria), to opportunistic human pathogens, to free-living species in soils, sediments and water (Carareto Alves et al. 2014). In this respect, the general term “agrobacteria” refers to a polyphyletic group of *Rhizobiaceae* that are able to cause neoplastic diseases on plants (de Lajudie et al. 2019).

Agrobacteria are remarkable plant pathogens, as the infection process represents an interkingdom genetic exchange involving integration of a fragment of bacterial plasmid DNA (transferred DNA or T-DNA) into plant host cells (Gelvin 2017). Consequently, agrobacteria cause crown and cane gall (Escobar and Dandekar 2003; Puławska 2010), and hairy root (Bosmans et al. 2017) diseases, depending on whether they carry a tumor-inducing (Ti) or root-inducing (Ri) plasmid. Hence, Ti and Ri plasmids code for functions essential for pathogenicity. Ti and Ri plasmids are related, although the former plasmid group has been studied more extensively.

Ti plasmids are transmissible and self-conjugal (reviewed in (Farrand 1998)). However, the natural host range of Ti plasmids is relatively narrow and restricted to members of the family *Rhizobiaceae*. To date, strains carrying Ti plasmids and able to cause crown gall and hairy root diseases (agrobacteria) have been primarily identified within the genera *Agrobacterium*, *Allorhizobium*, and *Rhizobium*. Additionally, a *Neorhizobium* strain carrying a Ti plasmid and able to cause tumors on multiple host plants was identified recently (Haryono et al. 2018). Historically, *Rhizobium rhizogenes* (i.e. *Agrobacterium* biovar 2/*Agrobacterium rhizogenes*) was the only tumorigenic *Rhizobium* species. However, another member of this genus, *Rhizobium tumorigenes*, was recently isolated from cane gall tumors on thornless blackberry (Kuzmanović et al. 2018). In addition, genomic analyses now suggest that the tumorigenic strain AB2/73 (Anderson and Moore 1979), initially identified as a biovar 2 strain (*R. rhizogenes*) (Unger et al. 1985), actually belongs to a novel, so far undescribed *Rhizobium* species (Hooykaas and Hooykaas 2021).

In our previous work, we identified a novel group of tumorigenic agrobacteria associated with crown gall disease of rhododendron (Kuzmanović et al. 2019). Phylogenetic and genomic analyses suggested that these strains are most closely related *R. tumorigenes*, but represent a separate species. Collectively, we named this distinct *Rhizobium* clade comprising *R. tumorigenes* and novel rhododendron strains as “tumorigenes”. In this study, we generated high quality, closed genomes of representatives of the “tumorigenes” clade, and performed thorough comparative genomic and phylogenomic analyses. Moreover, we phenotypically characterized rhododendron strains and described them as a novel species, *Rhizobium rhododendri*.

## 2 | MATERIALS AND METHODS

### 2.1 | Bacterial strains

*Rhizobium* strain rho-6.2^T^ (= DSM 110655^T^ = CFBP 9067^T^) used in this study was isolated in 2017 from crown gall tumors on rhododendron originating from a nursery in Lower Saxony, Germany (Kuzmanović DSM 104880^T^ = CFBP et al. 2019). In addition, we used *R. tumorigenes* strains 1078^T^ (= 8567^T^) and 932 (= DSM 104878 = CFBP 8566) reported in our previous study (Kuzmanović et al. 2018). For whole genome sequencing, bacteria were grown in tryptone-yeast (TY) broth (tryptone 5 g/l, yeast extract 3 g/l, CaCl_2_×2H_2_O 0.9 g/l) at 28°C for 48 h. Cultures were stored in a −80°C freezer in nutrient broth with 20% glycerol for long-term preservation.

### 2.2 | Phenotypic characterization and fatty acid methyl ester (FAME) analysis

The growth of bacterial strains rho-6.2^T^ and 1078^T^ was assessed on different agar media: yeast mannitol agar (YMA) (Kuzmanović et al. 2015), TY, R2A (DSMZ medium 830), potato dextrose agar supplemented with 0.08% CaCO_3_ (PDA-CaCO_3_) (Bouzar et al. 1995), and King’s medium B (King et al. 1954). Their motility was examined microscopically. The Gram reaction was determined by KOH (Ryu 1939) and aminopeptidase (Cerny 1976) (Bactident Aminopeptidase, Merck, Cat. No.113301, Germany) tests. Oxidase activity was tested by the method of Kovacs (1956). Catalase tests were performed by mixing freshly grown bacterial cells with 10% H_2_O_2_, followed by examination of gas bubble formation. Growth at 5, 10, 15, 20, 25, 30, 35, and 40°C was determined in R2A broth for up to 9 days. Tests for 3-ketolactose production, and acid clearing on PDA-CaCO_3_ were performed as described before (Moore et al. 2001). Additionally, the strains rho-6.2^T^ and 1078^T^ were phenotypically characterized using the API 20NE system (bioMérieux, Marcv LEtoile, France) following the instructions provided by the manufacturer.

For the fatty acid methyl esters (FAME) analysis, strains were cultured on R2A medium at 25°C for three days. The cellular fatty acids were analyzed using the Microbial Identification System (MIDI; Sherlock version 6.1, TSBA40 method), according to instructions provided by the manufacturer (Sasser 1990). A combined analysis by gas chromatography coupled to a mass spectrometer was used to confirm the identity of the fatty acids based on retention time and mass spectral data (Vieira et al. 2021).

### 2.3 | DNA extraction

Genomic DNA was extracted from bacterial strains using a Qiagen Genomic DNA Buffer Set (Qiagen, Germany; Cat. No. 19060) and Qiagen genomic tip 100/G gravity-flow, anion exchange columns (Cat. No. 10243). The purity and approximate concentration of DNA was determined by spectrophotometry using the NanoDrop instrument. Genomic DNA integrity was assessed by agarose gel electrophoresis.

### 2.4 | Eckhardt-type gel electrophoresis

Plasmid content of *Rhizobium* strains rho-6.2^T^, 1078^T^ and 932 was analyzed by the modified method of Eckhardt (1978). This method can also allow visualization of other extrachromosomal replicons, such as smaller chromids. Separation and visualization of replicons was performed in a 0.7% (w/v) agarose gel (5 mm thick) prepared in 1× Tris-borate-EDTA (TBE) buffer using the following procedure. Bacteria were grown in TY medium for 24 h at 28°C. Approximately 0.5-1 mL of bacterial culture was centrifuged at 8,000 rcf (g) for 10 min, and the pellet was resuspended in 0.5 mL sterile distilled water. One mL of 0.3% (m/v) sodium lauroylsarcosinate was added, after which the cell suspension was gently mixed and centrifuged at 8,000 rcf for 5 min. The pellet was resuspended in 40 µL 20% Ficoll 400 (w/v) in TE buffer (10 mmol L^−1^ Tris–HCl, 1 mmol L^−1^ EDTA, pH 8.0) and samples were incubated for 15 min on ice. An agarose gel was prepared during the previous incubation steps by loading 25 μL of 10% sodium dodecyl sulfate (SDS; w/v) into empty wells, followed by gently flooding the gel with 1x TBE buffer and running electrophoresis at 4 V cm^−1^ for 15 min from positive to negative polarity (opposite direction to standard DNA gel electrophoresis). Next, 10 µL lysing solution in TE buffer containing 0.4 mg mL^−1^ RNase A, 1 mg mL^−1^ bromophenol blue, and 1.5 mg mL^−1^ lysozyme (freshly prepared aqueous solution) was added to each cell sample after incubation on ice. A 30 µL aliquot of the mixture was loaded immediately into wells in the gel. Electrophoresis was run first at 1.5 V cm^−1^ for 1h and then at 4 V cm^−1^ for 20h (standard DNA gel electrophoresis from negative to positive polarity). The gel was stained in ethidium bromide solution (1 μg/mL) and the plasmids were visualized under UV light. As markers, “*Agrobacterium fabrum*” C58^T^, *Allorhizobium ampelinum* S4^T^, and *R. rhizogenes* K84 carrying replicons of known size were used.

### 2.5 | Illumina library preparation and sequencing

Libraries for Illumina sequencing were prepared using a Nextera XT DNA Library Preparation Kit (Illumina, San Diego, USA) with modifications according to Baym (Baym et al. 2015). Genome sequencing of strains 1078^T^ and 932 was performed using an Illumina NextSeq 500 platform in PE75 mode. For strain rho-6.2^T^, paired-end 151 bp reads previously generated on an Illumina NextSeq 500 platform (Kuzmanović et al. 2019) were used for error-correction of the PacBio assembly (see below).

### 2.6 | PacBio library preparation and sequencing

SMRTbell template libraries were prepared according to the instructions from Pacific Biosciences (Menlo Park, CA, USA), following the Procedure & Checklist – Greater Than 10 kb Template Preparation document. Briefly, for preparation of 15 kb libraries, 8 µg genomic DNA was sheared using g-tubes from Covaris (Woburn, MA, USA) according to the manufactureŕs instructions. DNA was end-repaired and ligated overnight to hairpin adapters applying components from the DNA/Polymerase Binding Kit P6 from Pacific Biosciences. Reactions were carried out according to the instructions of the manufacturer. BluePippin Size-Selection to greater than 4 kb was performed according to the manufactureŕs instructions (Sage Science, Beverly, MA, USA). Conditions for annealing of the sequencing primers and binding of polymerase to purified SMRTbell template were assessed with the Calculator in RS Remote (Pacific Biosciences). Single-molecule real-time (SMRT) sequencing was carried out on the PacBio RSII (PacificBiosciences) taking one 240-minutes movie on one SMRT cell per sample using the P6 Chemistry.

### 2.7 | Genome assembly, error-correction and annotation

SMRT Cell data were assembled using the “RS_HGAP_Assembly.3” protocol included in SMRT Portal version 2.3.0 using default parameters. Assembled replicons were circularized and adjusted to *dnaA* (chromosomes) or *repA* (chromids and megaplasmids) as the first gene.

Error-correction was performed by mapping Illumina paired-end reads (2×150 bp for rho-6.2^T^, and 2×75 bp for strains 932 and 1078^T^) onto the PacBio assemblies using BWA 0.6.2 (Li and Durbin 2009) with subsequent variant and consensus calling using VarScan 2.3.7 (Koboldt et al. 2012). Moreover, to visually inspect and manually correct the remaining errors, long and short reads were mapped to the assembled sequences with minimap2 (Galaxy Version 2.17+galaxy0) (Li 2018) and Bowtie2 (Galaxy Version 2.4.5+galaxy0) (Langmead and Salzberg 2012), respectively. Consensus concordances of QV60 were confirmed for all three genomes.

Finally, genome sequences were annotated. For all analyses reported in this study, annotations produced by Prokka (Galaxy Version 1.13) (Seemann 2014) were used. Annotation of particular sequences of interest and metabolic pathway prediction were performed using eggNOG-mapper (version emapper-2.1.9) (Cantalapiedra et al. 2021) based on eggNOG orthology data (Huerta-Cepas et al. 2018), as well as with BlastKOALA (last accessed on November, 2022) (Kanehisa et al. 2016). For eggNOG-mapper, sequence searches were performed using DIAMOND version 2.0.11 (Buchfink et al. 2021). Moreover, to aid functional annotation of some loci, BLASTp comparison against the NCBI non-redundant (nr) protein database (https://blast.ncbi.nlm.nih.gov/Blast.cgi; last accessed on November, 2022) (Johnson et al. 2008) was conducted. Prophage prediction was done using PHASTER web server (https://phaster.ca/; last accessed on November, 2022) (Arndt et al. 2016). Insertion sequence (IS) elements were identified using ISEscan version 1.7.2.3 (Xie and Tang 2017).

### 2.8 | Classification, synteny and phylogeny of DNA replicons

Bacterial replicons were classified using an approach similar to that described previously (diCenzo and Finan 2017; diCenzo et al. 2019), except that a size threshold was not used in defining megaplasmids or chromids (Hall et al. 2022). The largest replicon in a genome was classified as the chromosome. The remaining replicons were considered putative chromids if both their %GC content and dinucleotide relative abundance (DRA) distance differed by not more than approximately 1% and 0.4, respectively, compared to the chromosome. DRA distances were computed as described by diCenzo and Finan (2017). The replicons that failed to meet one of the two criteria (%GC-and DRA distance-based) for chromid classification were further analyzed by means of comparative genomic and phylogenetic analysis to reconstruct their evolutionary history, as described in the following paragraphs. The remaining replicons that did not meet any of above-mentioned criteria were classified as megaplasmids.

Synteny between genomes of the “tumorigenes” clade was explored using circos. First, blast bidirectional best hits (Blast-BBHs) were identified using BLASTn version 2.10.1+ (Camacho et al. 2009) and a custom Matlab script, limiting Blast-BBHs to those with pairs where at least 50% of each protein was aligned with e-values ≤1e-100. The parsed output was then used to prepare a “links” file, which was provided to circos 0.69-8 (Krzywinski et al. 2009). The scripts are available at https://github.com/diCenzo-Lab/007_2023_Rhizobium_rhododendri. Furthermore, BRIG (BLAST Ring Image Generator) program version 0.95 (Alikhan et al. 2011) was used for visual representation of replicons of strain rho-6.2^T^ (reference sequences) with the orthologous replicons of related strains (query sequences). The BRIG analysis was done by using the BLASTn option.

To assess the evolutionary relationships among the extrachromosomal replicons of the “tumorigenes” clade, phylogenetic analysis based on the RepA and RepB protein sequences was conducted. Protein sequence alignments for each set of orthologs were generated using MAFFT version 7 (Katoh et al. 2017). Maximum likelihood (ML) phylogenies based on individual RepA and RepB sequences and their concatenation were inferred using IQ-TREE 1.6.12 (Nguyen et al. 2015) available through the IQ-TREE web server (http://iqtree.cibiv.univie.ac.at/) (Trifinopoulos et al. 2016). Model selection was conducted using IQ-TREE ModelFinder (Kalyaanamoorthy et al. 2017) based on Bayesian Information Criterion (BIC) (Schwarz 1978). Branch supports were assessed by ultrafast bootstrap analysis (UFBoot) (Hoang et al. 2017) and the SH-aLRT test (Guindon et al. 2010) using 1000 replicates. The trees were visualized using FigTree version 1.4.4 (https://github.com/rambaut/figtree) and edited using Inkscape version 1.2.1 (https://inkscape.org/).

To examine potential relationships between the extrachromosomal replicons of the “tumorigenes” clade and other members of the family *Rhizobiaceae*, a previously described pipeline was adapted (diCenzo et al. 2019) and is available at https://github.com/diCenzo-Lab/007_2023_Rhizobium_rhododendri. Shortly, putative RepA proteins were identified in each of the *Rhizobiaceae* proteomes using the hmmsearch function of HMMER version 3.3 (Eddy 2009) and the Pfam ParA hidden Markov model (HMM). All hits were then searched against the complete Pfam version 34.0 and TIGERFAM version 15.0 databases (Finn et al. 2016; Haft et al. 2013), and proteins were classified as RepA if the top was either the ParA (Pfam) or TIGR03453 (TIGRFAM) HMM. All RepA proteins were aligned with MAFFT version 7.471 (Katoh and Standley 2013) with the ‘localpair’ option, and then trimmed with trimAl version 1.4.rev22 (Capella-Gutiérrez et al. 2009) with the ‘automated1’ option. A maximum likelihood phylogeny was constructed using RAxML version 8.2.12 (Stamatakis 2014) with the LG amino acid substitution model with empirical base frequencies and the final tree represents the bootstrap best tree following 500 bootstrap replicates. In addition, RepA proteins were clustered using CD-HIT version 4.8.1 (Li and Godzik 2006) with a 90% identity threshold.

### 2.9 | Genome-based phylogenetic analyses

The dataset comprised of 119 genomes, including 116 *Rhizobiaceae* strains and three *Mesorhizobium* spp. that were used as an outgroup (Table A1). In particular, finished genomes of *Rhizobium* strain rho-6.2^T^ and *R. tumorigenes* strains 1078^T^ and 932 obtained in this study were used. We also included the previously-reported draft genome sequences of two additional *Rhizobium* strains associated with rhododendron crown gall, rho-1.1 and rho-13.1 (Kuzmanović et al. 2019). Genome sequences of related *Rhizobium* strains, as well as representatives of various *Rhizobiaceae* genera, were retrieved from GenBank. The genomes closely related to the “tumorigenes” clade representatives were identified by NCBI BLASTn (https://blast.ncbi.nlm.nih.gov/Blast.cgi) searches against the nucleotide collection (nr/nt) and whole-genome shotgun contigs (wgs) databases using 16S rRNA and *recA* housekeeping gene sequences as a query, with default parameters (last accessed on November, 2022).

Core-genome- and pan-genome-based phylogenies were inferred using GET_HOMOLOGUES Version 11042019 (Contreras-Moreira and Vinuesa 2013) and GET_PHYLOMARKERS Version 2.2.8_18Nov2018 (Vinuesa et al. 2018) as described before (Kuzmanović et al. 2022b). For core-genome-based phylogenetic analyses, the latter pipeline was run using both DNA and protein sequences, thus generating core-genome and core-proteome phylogenies, respectively. The protein alignment generated by the cpAAI pipeline (see below) was also used as input for phylogenetic analysis. A ML phylogeny was inferred under the best-fitting substitution model by employing IQ-TREE Version 2.1.3 (Nguyen et al. 2015) and ModelFinder (integrated in IQ-TREE) (Kalyaanamoorthy et al. 2017), following the same approach as implemented in the GET_PHYLOMARKERS package.

### 2.10 | Genome and proteome relatedness indices

For calculation of genome and proteome relatedness indices, we used the same dataset as for phylogenetic analysis (see above; Table A1). For delineation of species, we computed overall genome relatedness indices (OGRIs), in particular, average nucleotide identity (ANI) (Goris et al. 2007; Richter and Rossello-Mora 2009) and digital DNA-DNA hybridization (dDDH) (Meier-Kolthoff et al. 2013). The ANI calculations were performed using PyANI Version 0.2.11, with scripts employing BLAST+ (ANIb) to align the input sequences (https://github.com/widdowquinn/pyani) (Pritchard et al. 2016), OrthoANIu Version 1.2 (calculates orthologous ANI using USEARCH algorithm) (Yoon et al. 2017), and FastANI Version 1.2 (estimates ANI using Mashmap as its MinHash-based alignment-free sequence mapping engine) (Jain et al. 2018). The dDDH values were computed by the Genome-to-Genome Distance Calculator (GGDC 3.0) implemented in the Type (Strain) Genome Server (TYGS) (Meier-Kolthoff et al. 2021; Meier-Kolthoff and Göker 2019). The dDDH values calculated under the formula 2 (GBDP formula d_4_: identities/HSP length) were considered (Meier-Kolthoff et al. 2013).

For delineation of genera, we computed whole-proteome average amino-acid identity (wpAAI; more commonly known as AAI) (Goris et al. 2007; Konstantinidis et al. 2017; Konstantinidis and Tiedje 2005) and core-proteome average amino-acid identity (cpAAI) (Kuzmanović et al. 2022a). The wpAAI values were computed using the CompareM software (github.com/dparks1134/CompareM) using the aai_wf command with default parameters. For calculation of cpAAI, the cpAAI_Rhizobiaceae pipeline (github.com/flass/cpAAI_Rhizobiaceae) was employed to generate a concatenated protein alignment of a reference set of 170 marker proteins from 97 reference strains, using the pre-aligned reference protein files (option –A) as described in Kuzmanović et al. (Kuzmanović et al. 2022a). Nucleotide FASTA files of all CDSs predicted by Prokka were used as input files. The cpAAI values were computed from the resulting alignment using a custom R script (see github.com/flass/cpAAI_Rhizobiaceae) that relied on the “dist.aa” function from the “ape” package (Paradis and Schliep 2018). Additionally, we calculated cpAAI values using the core protein markers inferred from the 119 strains included in the present study, which were identified and selected using the GET_HOMOLOGUES and GET_PHYLOMARKERS tools, respectively (see above).

Heatmaps representing genome (OGRIs) and proteome (cpAAI and wpAAI) relatedness values were generated and plotted onto the reference core-proteome phylogenetic tree by the “phylo.heatmap” function in the R package phytools (Revell 2012).

### 2.11 | Identification of species-specific genes

In order to identify genes specific for each of the two species comprising the “tumorigenes” clade, the pan-genome of the “tumorigenes” clade was explored. The dataset included two *R. tumorigenes* strains (1078^T^ and 932) and 12 strains comprising the new species *Rhizobium rhododendri* (see below) (Table A1). The analysis was performed using the GET_HOMOLOGUES software and its auxiliary scripts as described before (Kuzmanović et al. 2020).

To determine if the function of the species-specific gene or gene cluster of interest is compensated by isoenzymes or by a divergent homologous gene(s) in the other species, we performed BLASTp (Johnson et al. 2008) comparisons, and examined annotations of pan-genome genes of species comprising the “tumorigenes” clade that were generated by GhostKOALA (Kanehisa et al. 2016) and eggNOG-mapper (Cantalapiedra et al. 2021).

## 3 | RESULTS

### 3.1 | Genome sequences and rRNA operon diversity

The finished genome sequences of three representative members of the *Rhizobium* clade “tumorigenes” (rho-6.2^T^, 1078^T^ and 932) were generated using a combination of long- (PacBio) and short-read (Illumina) sequencing technologies (see Table A2 for summary statistics of the generated sequencing data). Genome assembly and polishing resulted in gapless, circular replicons for all sequenced strains, with high average sequencing depths (>200×, >140× for long read data) (Table 2). The genome of strain rho-6.2^T^ was composed of four replicons, while six replicons were identified in each of the strains 1078^T^ and 932. The presence of smaller replicons (approximately <1.5 Mb) was confirmed by a modified Eckhardt agarose gel electrophoresis technique, although some of the replicons of similar size could not be clearly differentiated (Figure A1). The total genome size of the three strains was similar, ranging from 5.96 to 5.98 Mb (Table 1). The GC content was approximately 60% for all strains (Table 1).

**Table 1.** General features of the genome sequences obtained in this study.

|  | Strains |  |  |
| --- | --- | --- | --- |
|  | *Rhizobium rhododendri*<br>rho-6.2<sup>T</sup> | *Rhizobium tumorigenes*<br>1078<sup>T</sup> | *Rhizobium tumorigenes*<br>932 |
| Replicons | 4 | 6 | 6 |
| Size (Mb) | 5.96 | 5.98 | 5.97 |
| GC content (%) | 59.98 | 59.96 | 60.03 |
| Genes<sup>a</sup> | 5,649 | 5,685 | 5,705 |
| CDSs<sup>a</sup> | 5,582 | 5,619 | 5,637 |
| rRNA operons (5S, 23S, 16S) | 4 | 4 | 4 |
| Genome coverage | 209× | 329× | 285× |

<sup>a</sup>Numbers based on Prokka annotation.

**Table 2.** Classification of replicons and their general features. All replicons were circular.

| Replicon | Size (bp) | GC% | Accession<br>Number<sup>a</sup> |
| --- | --- | --- | --- |
| ***Rhizobium rhododendri* rho-6.2<sup>T</sup>** |  |  |  |
| Chromosome | 3,709,686 | 60.92 | X |
| Putative chromid 1 | 1,530,638 | 58.41 | X |
| Megaplasmid pTi6.2 | 381,845 | 56.14 | X |
| Putative chromid 2 | 336,962 | 60.98 | X |
| ***Rhizobium tumorigenes* 1078<sup>T</sup>** |  |  |  |
| Chromosome | 3,664,408 | 60.77 | X |
| Megaplasmid pRt1078 | 834,411 | 57.79 | X |
| Megaplasmid pTi1078 | 439,071 | 56.21 | X |
| Putative chromid 1 | 432,998 | 60.2 | X |
| Putative chromid 2 | 304,572 | 61.07 | X |
| Putative chromid 3 | 302,267 | 60.24 | X |
| ***Rhizobium tumorigenes* 932** |  |  |  |
| Chromosome | 3,816,680 | 60.66 | X |
| Megaplasmid pRt932 | 756,443 | 58.29 | X |
| Megaplasmid pTi932 | 430,508 | 56.05 | X |
| Putative chromid 1 | 339,618 | 60.61 | X |
| Putative chromid 2 | 319,653 | 60.41 | X |
| Putative chromid 3 | 304,177 | 61.03 | X |

<sup>a</sup>NCBI submission is undergoing processing and accession numbers will be added when available.

For all three strains, four rRNA operons (5S, 16S, and 23S rRNA) were identified on the largest replicon. Unlike strains rho-6.2^T^ and 1078^T^, we did not observe intragenomic heterogeneity between multiple rRNA operons in strain 932. In strain rho-6.2^T^, one of the variants of the rRNA operon differed by only one SNP in the 5S rRNA gene from the remaining three copies. For strain 1078^T^, we identified three different variants of the rRNA operons. The first variant encompassed two rRNA copies and differed by one SNP from the second variant, whereas several INDELs and SNPs were identified when compared to the third variant. The sequence variations were located in the 23S rRNA gene and 16S-23S ITS region. The 16S rRNA gene sequences were identical across both *R. tumorigenes* strains (1078^T^ and 932), and differed by 10 SNPs from those of strain rho-6.2^T^.

### 3.2 | Genome organization

Whole-genome sequencing revealed that all strains in the “tumorigenes” clade contain multipartite genomes (Figure 1; Table 2). The largest replicon in all sequenced genomes, which also carried all four copies of the rRNA operon, was classified as the chromosome. Chromosomes were highly conserved across all strains of the “tumorigenes” clade (Figure 2; Figure A2a).

**Figure 1.**
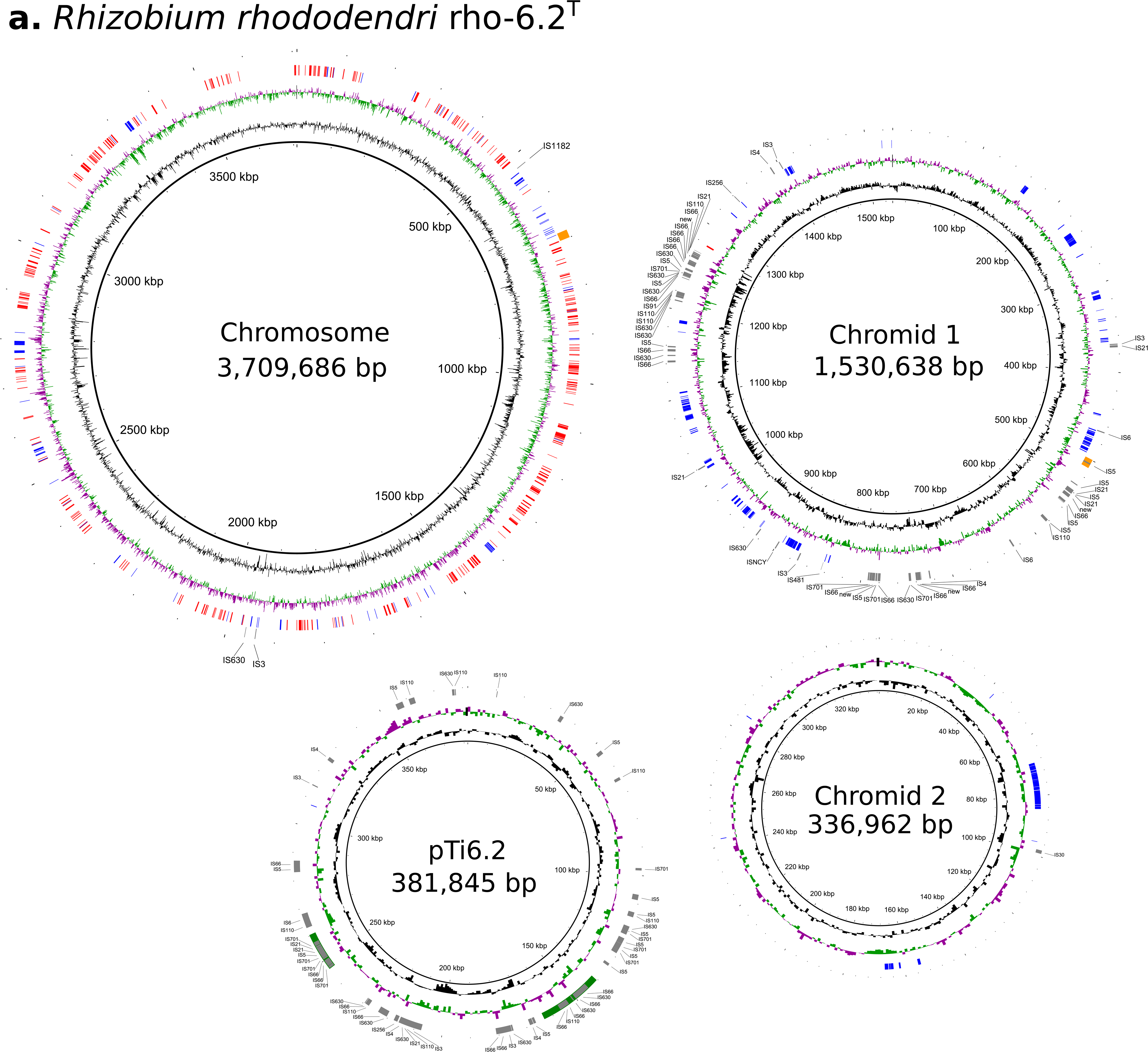
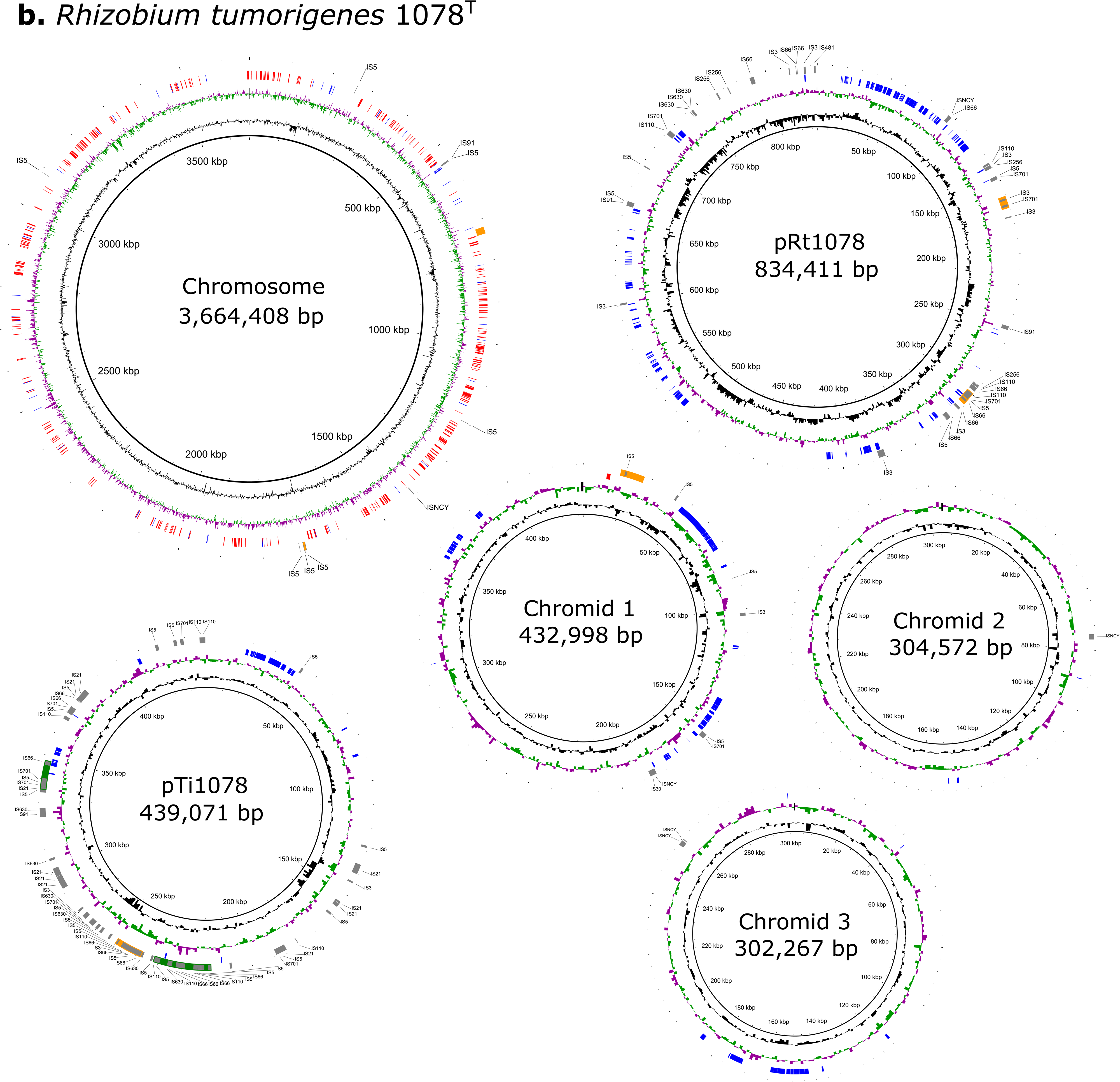
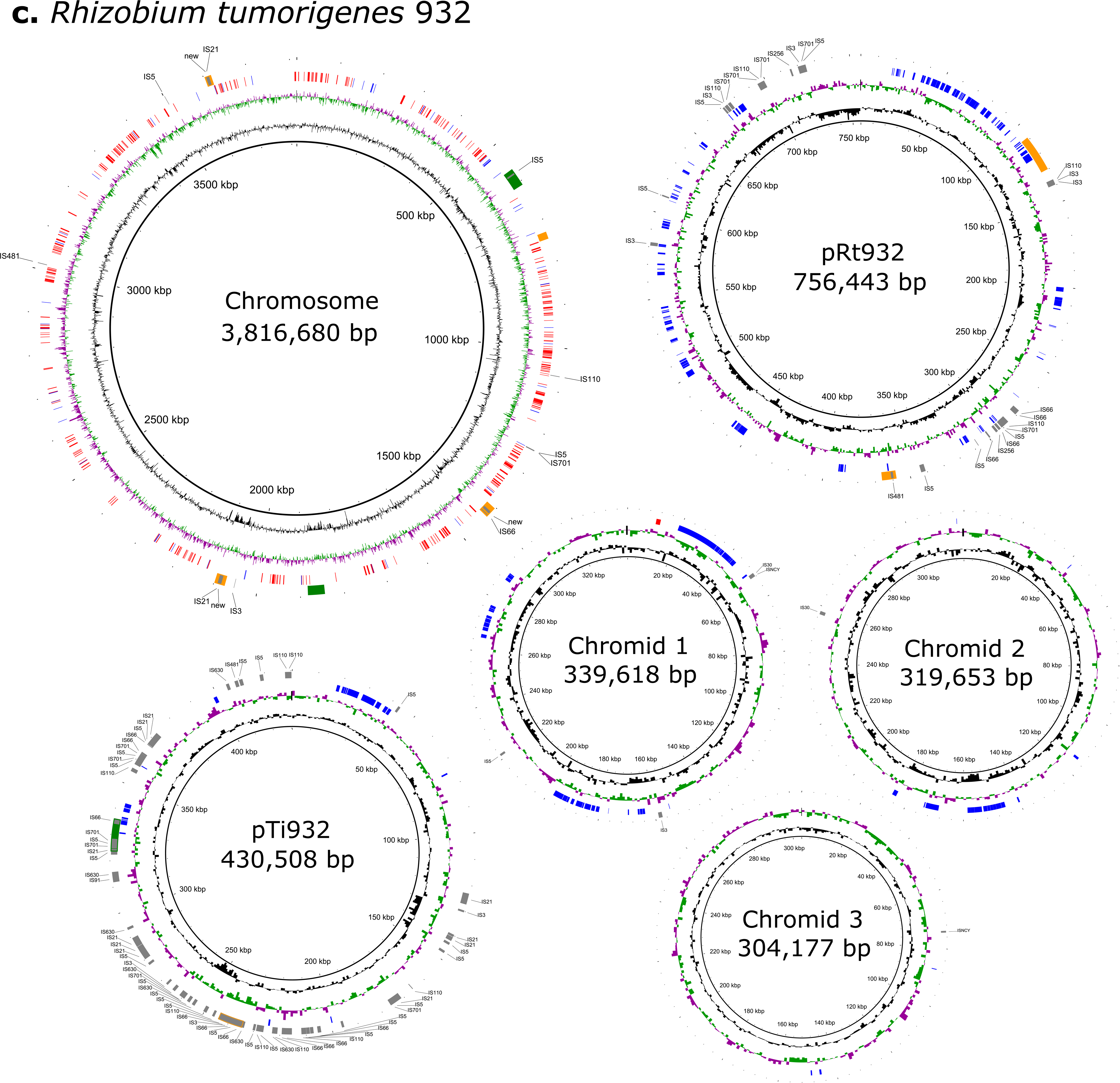
Circular maps of the complete genomes of strain *Rhizobium rhododendri* rho-6.2^T^ (a), and strains of *Rhizobium tumorigenes* 1078^T^ (b) and 932 (c). Each replicon is presented by a circular plot containing five rings. Genetic coordinates of the reference sequences are shown within the thin inner ring. The next two rings portray GC content (black ring) and GC skew (purple/green). The next ring shows core (red) and species-specific (blue) genes. Core genes (364) were identified from a dataset of 119 strains using GET_HOMOLOGUES software. The accessory genes (*R. rhododendri* vs. *R. tumorigenes*) were identified with the same software. The outermost ring highlights prophage regions identified with PHASTER (intact prophages are shown in green and incomplete in orange) and insertion sequence (IS) elements identified using ISEscan (shown in gray). As in some cases IS elements were identified within the prophage regions, borders of the latter regions are highlighted with the corresponding color. The Figure was generated using BRIG software and edited with Inkscape (see M&M for details).

**Figure 2.**
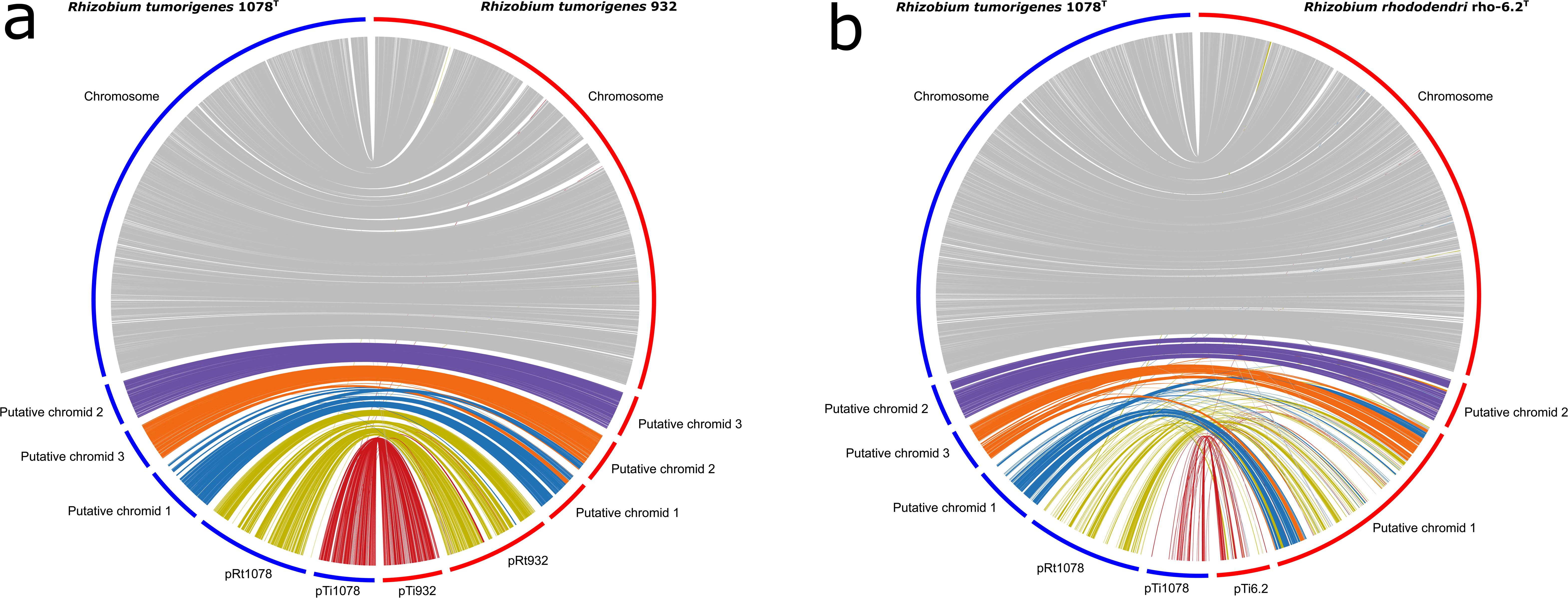
Synteny analysis of the genomes of the “tumorigenes” clade representatives. The *Rhizobium tumorigenes* 1078^T^ genome was compared with the genome of *R. tumorigenes* 932 (a) and *Rhizobium rhododendri* rho-6.2^T^ (b). Putative orthologous genes between strains were identified by performing BLAST bidirectional best-hit analyses using the proteomes. BLAST 1 × 10−100 and ≥ corresponding gene, and their position was mapped on the genome. Each putative ortholog between genomes is connected by a line and color coded based on the location of the gene in the *R. tumorigenes* 1078^T^ genome. The Figure was generated using circos software and edited with Inkscape (see M&M for details).

In accordance with our previous work demonstrating the pathogenicity of the “tumorigenes” clade (Kuzmanović et al. 2019; Kuzmanović et al. 2018), all three strains harbored a Ti megaplasmid. In addition, each of *R. tumorigenes* strains 1078^T^ and 932 carried an additional megaplasmid (756-835 kb) and three putative chromids (302 to 433 kb) (Tables 2 and A3), whose gene contents were highly conserved between strains 1078^T^ and 932 (Figure 2a). Interestingly, we also detected evidence of DNA exchange between replicons. In particular, a 41-gene cluster of putative chromid 1 of strain 1078^T^ (orthologous to putative chromid 1 of strain 932) was found on putative chromid 2 of strain 932 (orthologous to putative chromid 3 of strain 1078^T^). Likewise, a 25-gene cluster of putative chromid 3 of strain 1078 was found on putative chromid 1 of strain 932 (Figure 2a). Based on the location of these gene clusters in the more distantly related strain rho-6.2^T^ (Figure 2b), the observed translocations likely occurred in the lineage leading to strain 1078^T^ following divergence from strain 932.

Unlike *R. tumorigenes* strains 1078^T^ and 932 that carried five extrachromosomal replicons, strain rho-6.2^T^ carried only three: a Ti plasmid and two putative chromids. Surprisingly, synteny analysis suggested that putative chromid 1 of rho-6.2^T^ resulted from the cointegration of an ancestral megaplasmid (orthologous to pRt1078 of strain 1078^T^) and two putative chromids (orthologous to putative chromids 1 and 3 of 1078^T^) (Figure 2b). Consistent with the proposed cointegration scenario, the cointegrant of strain rho-6.2^T^ (putative chromid 1) contains three *repABC* cassettes, which are orthologous to the *repABC* cassettes of pRt1078 and putative chromids 1 and 3 of strain 1078^T^ (Figure A3). However, only one *repC* copy on the cointegrant is complete, with the other two copies appearing to be truncated and thus non-functional. The orthologous cointegrant replicon was also carried by other *R. rhododendri* strains, showing a high degree of synteny (Figure A2b).

Putative chromid 2 of rho-6.2^T^ displayed high conservation with putative chromid 2 of strain 1078^T^ (Figure A2c), whereas poor conservation was observed when comparing the Ti plasmids of these strains (Figure 2b). Orthologous replicons of putative chromid 2 are also present in other *R. rhododendri* strains, with all exhibiting a high degree of synteny (Figure A2c). On the other hand, the RepA proteins of each of the three *R. tumorigenes* chromids formed their own cluster in the RepA phylogeny (Figure A4) and they shared less than 92% identity with all other RepA protein from the family *Rhizobiaceae*. Comparison of the *R. tumorigenes* chromids with the most closely related replicons from other *Rhizobiaceae* species using D-Geneies (Cabanettes and Klopp 2018) identified no obvious stretches of synteny. Overall, these results suggest that the three chromids of *R. tumorigenes* are specific to the “tumorigenes” clade.

We could not identify genes associated with mobilization or conjugation on the chromids of strains 1078^T^ and 932. On the other hand, megaplasmids pRt932 and pRt1078 carried gene clusters involved in conjugative transfer, including genes coding for conjugative relaxase (*traA*/*virD2*), coupling protein (*traG*/*virD4*), and T4SS proteins (VirB/Trb). Likewise, the large cointegrant of rho-6.2^T^ carries genes for conjugation. Interestingly, however, these genes were divergent to those carried on pRt932 and pRt1078. For instance, the VirB4 protein sequences of pRt1078 and the cointegrant of rho-6.2’s shared only 40.8% identity.

### 3.3 | Phylogeny of the clade “tumorigenes”

The core genome of the 119 strains included in the analysis was identified using GET_HOMOLOGUES and comprised 364 homologous gene clusters. Phylogeny was inferred from 253 DNA and 191 protein markers that were selected using the GET_PHYLOMARKERS software. Moreover, we also inferred phylogeny from the protein alignment outputted by the cpAAI pipeline, which represented the concatenated sequence of a reference set of 169 protein markers. Although the original dataset included 170 protein markers, one marker gene was missing in *Onobrychidicola muellerharveyae* TH2^T^, and we therefore excluded this marker from the analysis.

All the resulting phylogenies were highly congruent, showing almost identical phylogenetic relationships between *Rhizobiaceae* genera and major *Rhizobiaceae* clades (data not shown). The only difference was the position of the genus *Xaviernesmea*, which was an outgroup of the clade containing the genera *Ensifer*, *Pararhizobium*, and *Sinorhizobium* in the DNA-based phylogenetic tree, while it was grouped with *Pararhizobium* spp. in the protein-based phylogenetic trees. Additionally, the phylogenetic position of several taxa within some sub-clades differed slightly between trees. Regardless, the phylogenetic positions of the taxa that are the subject of this work were identical across trees, and we therefore show only the core-proteome phylogenetic tree based on 191 protein markers (Figures 3 and A5). *R. tumorigenes* (1078^T^ and 932) and strains isolated from rhododendron in Germany (rho-6.2^T^, rho-1.1, and rho-13.1) clustered within two sister sub-clades in the clade we previously defined as “tumorigenes” (Kuzmanović et al. 2019) (Figures 3 and A5). The sub-clade containing the three rhododendron strains also included nine other *Rhizobium* strains whose genomes were available in GenBank. The rhododendron clade could be further divided into two clusters. The first cluster comprised our three rhododendron strains and strain L51/94 isolated from blueberry in Oregon (USA), while the second clade consisted of eight *Rhizobium* strains isolated from Himalayan blackberry in Oregon (Weisberg et al. 2022). The “tumorigenes” clade falls within the so-called core *Rhizobium* species complex, although it was distantly related to other *Rhizobium* species. *R. tubonense* was the closest relative of the “tumorigenes” representatives, while other *Rhizobium* species grouped within the “tropici-rhizogenes” clade and the more distantly related “leguminosarum-etli” clade (Figures 3 and A5).

**Figure 3.**
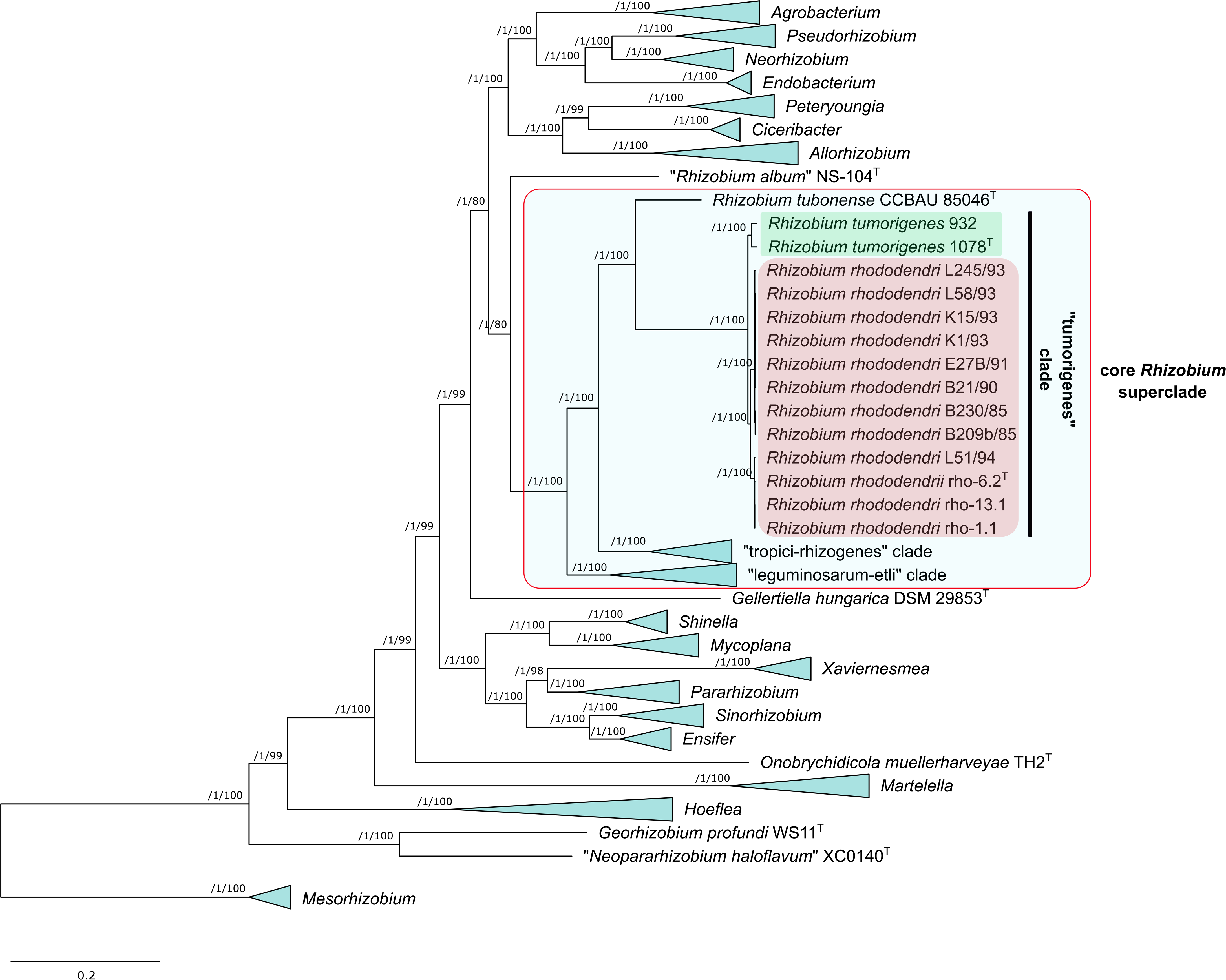
Maximum-likelihood core-proteome phylogeny showing the evolutionary relationships between and within the clade “tumorigenes” and other *Rhizobiaceae* clades (part collapsed). Three *Mesorhizobium* spp. strains were included as the outgroup to root the tree. The phylogeny was estimated from the concatenated alignments of 191 protein sequences selected as top-scoring markers using the GET_PHYLOMARKERS software. The numbers on the nodes indicate the approximate Bayesian posterior probabilities support values (first value) and ultra-fast bootstrap values (second value), as implemented in IQ-TREE. The scale bar represents the number of expected substitutions per site under the best-fitting LG+F+R6 model. The same tree, but without collapsing clades, is presented in the Figure A5.

A ML pan-genome phylogeny was estimated from a presence/absence matrix of 71,538 orthologous gene clusters. All *Rhizobiaceae* genera and major clades were resolved on the resulting tree (Figures 4 and A6), although their phylogenetic relationships differed from that determined from the core-proteome phylogeny (Figures 3 and A5). Nevertheless, the pan-genome phylogeny also contained the same two sub-clades within the “tumorigenes” clade: one comprising *R. tumorigenes*, and another with the rhododendron strains and those whose genomes were retrieved from the GenBank.

**Figure 4.**
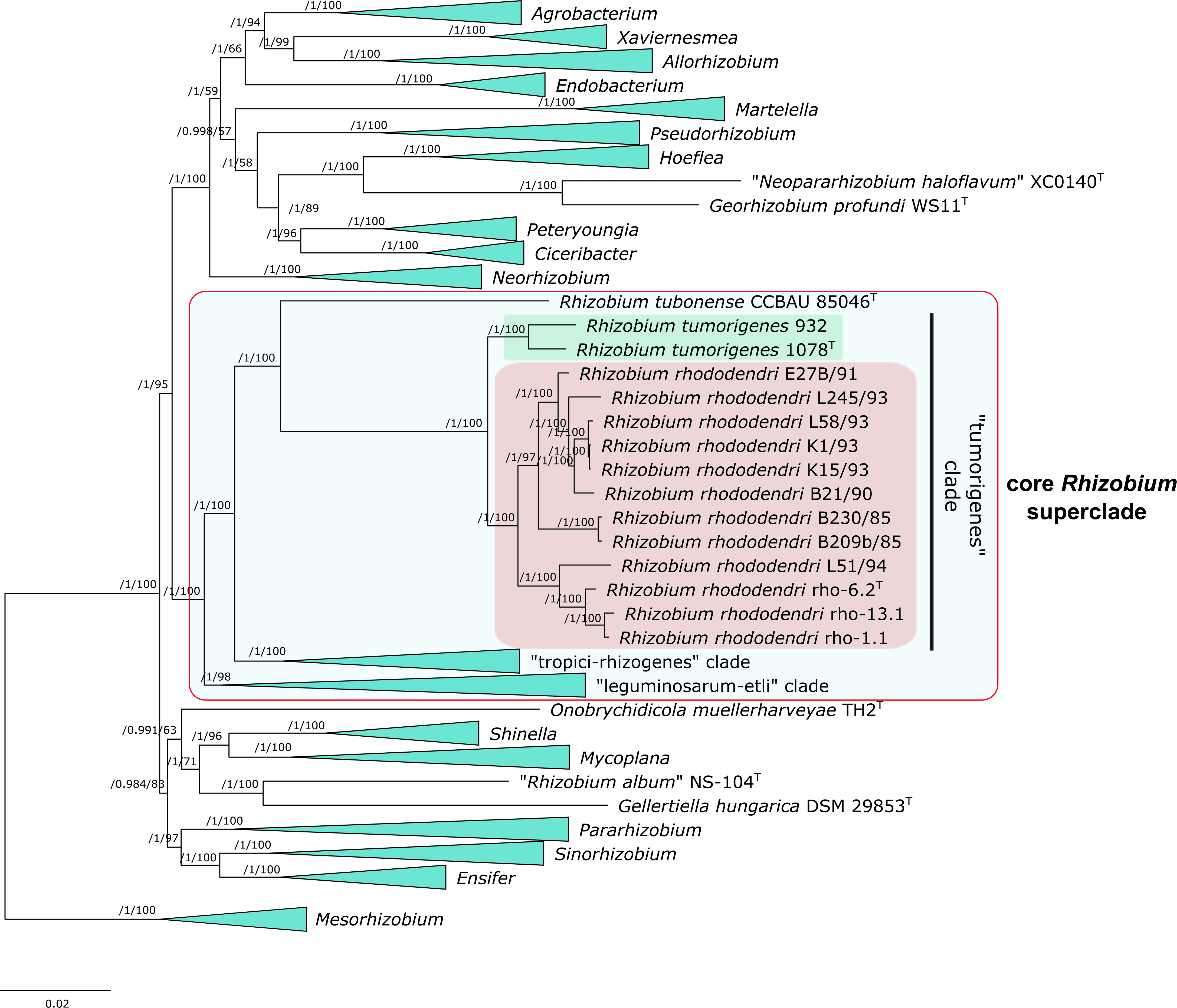
Maximum-likelihood pan-genome phylogeny showing the relationships between and within the clade “tumorigenes” and other *Rhizobiaceae* clades (part collapsed). Three *Mesorhizobium* spp. strains were included as the outgroup to root the tree. The tree was estimated with IQ-TREE from the consensus (COGtriangles and OMCL clusters) gene presence/absence matrix containing 71,538 clusters obtained using GET_HOMOLOGUES software. The numbers on the nodes indicate the approximate Bayesian posterior probabilities support values (first value) and ultra-fast bootstrap values (second value), as implemented in IQ-TREE. The scale bar represents the number of expected substitutions per site under the best-fitting GTR2+FO+R8 model. The same tree, but without collapsing clades, is presented in the Figure A65.

### 3.4 | Species delineation

For species delineation, we relied on ANI and dDDH computations. The threshold for species delineation was set at ~95-96% for ANI (Richter and Rossello-Mora 2009), consistent with previous recommendations (Goris et al. 2007). As for the conventional version of DDH, the generally accepted species boundary for dDDH values is 70% (Meier-Kolthoff et al. 2013; Stackebrandt and Goebel 1994). In this study, delineations of strains achieved by ANIb, OrthoANIu, FastANI, and dDDH were highly congruent. Several differences observed are discussed below. In any case, the OGRIs were consistent for the strains of the “tumorigenes” clade, which are the primary subject of this study (Table A4, Figures A7 and A8). The sub-clade containing *Rhizobium* strains isolated from rhododendron (strains rho-6.2^T^, rho-1.1 and rho-13.1) (Kuzmanović et al. 2019), blueberry (L51/94), and Himalayan blackberry (B21/90, B209b/85, B230/85, E27B/91, K1/93, K15/93, L51/94, L58/93 and L245/93) (Weisberg et al. 2022), and the sub-clade comprising species *R. tumorigenes* (1078^T^ and 932) (Kuzmanović et al. 2018) were clearly separated (Figures A7 and A8). When comparing members within the former sub-clade, they showed OGRIs >96.8% for ANIb and >75.1% for dDDH, indicating that they belong to the single species. In contrast, when compared to each other, the two sub-clades showed values <94.4% for ANIb and <57.3% for dDDH (Table A4). These results suggest that the two “tumorigenes” sub-clades represent distinct species, and we propose the name *Rhizobium rhododendri* (see the protologue below) for the sub-clade containing the strains originating from rhododendron, blueberry and Himalayan blackberry. During the writing of this manuscript, genomes of eight additional strains (VS19-DR96, VS19-DR104.1, VS19-DR104.2, VS19-DR121, VS19-DR129.2, VS19-DR181, VS19-DR183, and VS19-DRK62.2) became available in GenBank (BioProject Accession No. PRJNA762915) that also belong to the species *R. rhododendri*, based on ANIb comparisons (>99% ANI with rho-6.2^T^). These strains were isolated from cane galls of blueberry in Oregon (USA) in 2019. However, as these genomes were unpublished, they were not included in further analyses.

Phenotypic characteristics of strains *R. rhododendri* rho-6.2^T^ and *R. tumorigenes* 1078^T^ are listed in Table A5. As expected, these two strains showed almost identical phenotypic characteristics, and we were unable to identify clear differential characteristics. For the strain rho-6.2^T^, phenotypic characteristics are summarized in the protologue for the new species *R. rhododendri* (see below).

The results of the fatty acid analysis are summarized in Table A6. Similar to the other phenotypic characteristics that were measured, strains *R. rhododendri* rho-6.2^T^ and *R. tumorigenes* 1078^T^ exhibited highly similar FAME profiles. The only notable difference was in C_18:1_ w7c 11-methyl, which was ~2.5-fold more abundant in rho-6.2^T^ than in 1078^T^. Overall, the major fatty acids (>5%) identified in each of these strains are C_18:1_ ω7c (~50%), C_19:0_ cyclo ω7c (~18-22%), and C_16:0_ (~5-7%).

Moreover, our results suggested that *Rhizobium anhuiense* CCBAU 23252^T^ and *Rhizobium sophoriradicis* CCBAU 03470^T^ represent the same species (Table A4, Figures A7 and A8). Likewise, the following pairs of strains represent the same species based on our results (Table A4, Figures A7 and A8): *Rhizobium indigoferae* CCBAU 71042^T^ and *Rhizobium leguminosarum* USDA 2370^T^, *Rhizobium aethiopicum* HBR26^T^ and *Rhizobium aegyptiacum* 950, and *Rhizobium pisi* DSM 30132^T^ and *Rhizobium yanglingense* LMG 19592^T^. Additionally, ANIb and orthoANIu comparisons suggested that *Rhizobium favelukesii* LPU83^T^ and *Rhizobium tibeticum* CCBAU 85039^T^ also belong to the same species, although pairwise fastANI and dDDH values were at or slightly below the threshold for species delineation, respectively. For strains *Rhizobium dioscoreae* S-93^T^ and *Rhizobium* sp. AB2/73, and for *Rhizobium changzhiense* WYCCWR 11279^T^ and *Rhizobium sophorae* CCBAU 03386^T^, ANI values fell within the threshold of ~95-96%, while the dDDH values were slightly below the threshold of 70% (Table A4, Figures A7 and A8).

Furthermore, computed OGRIs suggested the existence of two new *Rhizobium* species within the clade “tropici-rhizogenes”. The first species comprised strains AC27/96 and Y79/96 isolated from Japanese pieris and rhododendron, respectively, and that were not designated as tumor inducing (Weisberg et al. 2020). The second potential new species included strains 17-2069-2b and 17-2069-2c isolated from blackberry, which were reported to carry Ti plasmids (Weisberg et al. 2022).

### 3.5 | Genus demarcation

Differentiation of *Rhizobiaceae* strains at the genus level was conducted using cpAAI and wpAAI indices (Table A7). Primarily, we relied on cpAAI calculated on the marker proteins selected in our former work (Kuzmanović et al. 2022a). As noted above, one marker was missing in *O. muellerharveyae* TH2^T^, and thus the comparison was based on 169 marker proteins. We used a cpAAI threshold of ~86%, combined with the core-proteome phylogeny shown in Figures 3 and A5, in considering genus delineation. As expected, genus demarcations (Figure 5) were generally consistent with our previous study (Kuzmanović et al. 2022a). However, the present dataset included a larger number of strains belonging to the core *Rhizobium* superclade compared to our previous analysis. Consistent with the core-proteome phylogeny, *Rhizobium* clades “tropici-rhizogenes”, “leguminosarum-etli” and “tumorigenes” were differentiated using a cpAAI threshold of 86%. Accordingly, these clades represent candidates for new *Rhizobiaceae* genera. However, delineation of *R. tubonense* was less clear. *R. tubonense* exhibited cpAAI values >86% with strains from both “tumorigenes” (86.79-86.93%) and “tropici-rhizogenes” (86.81-87.51%) clades, although this taxon was phylogenetically more closely related to the former clade. The wpAAI-based approach suggested the same unclear delineation of *R. tubonense* (Figure A9). On the other hand, based on cpAAI computed from 191 marker proteins selected in this study, *R. tubonense* exhibited cpAAI values slightly below 86% with “tumorigenes” and some “tropici-rhizogenes” clade members (Figure A10). For other *Rhizobiaceae* genera, all three methods (two cpAAI and wpAAI) were highly congruent, with a few differences observed within the genera *Hoeflea*, *Martelella*, and *Pararhizobium* (Figures 5, A9 and A10).

**Figure 5.**
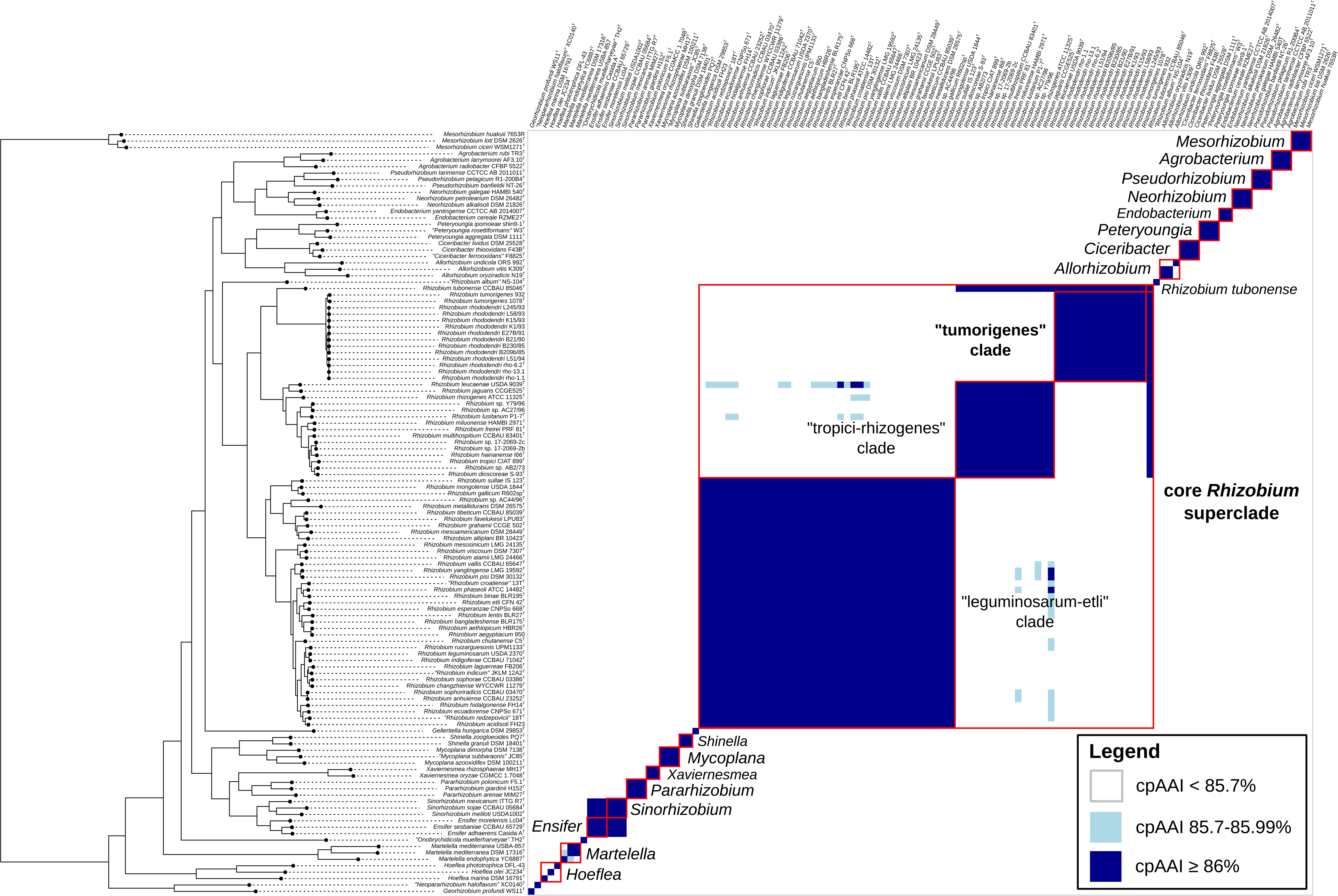
Clustered heatmap of core-proteome average amino-acid identity (cpAAI) values between the members of the clade “tumorigenes” and other *Rhizobiaceae* clades. Three *Mesorhizobium* spp. strains were included as the outgroup. cpAAI values were computed from a reference set of 169 protein markers defined in our previous study (Kuzmanović et al. 2022a). Although the original dataset included 170 protein markers, one marker gene was missing in *Onobrychidicola muellerharveyae* TH2^T^, and we therefore excluded this marker from the analysis. cpAAI values were clustered using the core-proteome phylogeny of Figures 3 and S5. The “tumorigenes” clade and other *Rhizobiaceae* clades are indicated with red boxes.

### 3.6 | Species-specific genes

#### 3.6.1 | R. rhododendri

Based on pan-genome analysis, 272 genes specific to *R. rhododendri* (*Rr*-specific) were identified. These genes were present in all 12 strains of *R. rhododendri* and absent in both *R. tumorigenes* strains. More than half (138) of these genes were located on putative chromid 1. Of the remaining genes, 108 were located on the chromosome, 27 on putative chromid 2, and 1 on pTi6.2 (Figure 1a). Most of the *Rr*-specific genes were annotated as hypothetical proteins or their function could not be clearly determined (Table A8a). Among those that were functionally annotated, based on COG categories, the most represented functional categories were K (Transcription), G (Carbohydrate metabolism and transport) and M (Cell wall/membrane/envelop biogenesis), comprising 32, 31, and 15 genes, respectively (Table A8a). Among the *Rr*-specific genes or gene clusters with predicted biological functions was the gene cluster Rr62_02696-Rr62_02698, predicted to be involved in production of cellulose (Table A8a); however, homologous, but divergent gene clusters with the same predicted function were also present in both *R. tumorigenes* strains (At1078_ 03513-At1078_ 03517 in strain 1078^T^), and in another copy also in strain rho-6.2^T^ (Rr62_03520-Rr62_03524). All putative gene clusters putatively associated with cellulose synthesis were located on chromosomes. Furthermore, the *Rr*-specific gene clusters Rr62_04000-Rr62_04012 and Rr62_05395-Rr62_05405 are annotated as being involved in the processing of various simple sugars (D-psicose/D-tagatose/L-ribulose) and sugar alcohols (galactitol, glucitol/sorbitol), respectively (Table A8a).

#### 3.6.2 | R. tumorigenes

Exploration of the pan-genome of the “tumorigenes” clade resulted in 322 genes that are specific to *R. tumorigenes* (*Rt*-specific), meaning they are present in both *R. tumorigenes* strains and absent from all *R. rhododendri* strains. In strain 1078^T^, one of these 322 genes was present in three copies, while there were two copies of five other genes. In strain 932, nine genes were present in two copies. Of the 326 genes specific for *R. tumorigenes* strain 1078^T^, including genes present in multiple copies, 159 were located on pRt1078, 58 on the chromosome, 57 on putative chromid 1, 31 on pTi1078, 3 on putative chromid 2, and 21 on putative chromid 3 (Figure 1b). Of the 331 genes in strain 932, 164 are located on pRt932, 59 on the chromosome, 54 on putative chromid 1, 30 on pTi1078, 21 on putative chromid 2, and 3 on putative chromid 3 (Figure 1c). Differences in the number of *Rt*-specific genes on each replicon may be explained by inter-replicon rearrangements as described above (see subsection “Genome organization”). As for the *Rr*-specific genes, the function of the majority of the *Rt*-specific genes could not be precisely determined (Table A8b). Based on COG classification, annotated *Rt*-specific genes of strain 1078^T^ were primarily annotated as belonging to the functional categories P (Inorganic ion transport and metabolism; 25 genes), L (Replication and repair; 24 genes), K (Transcription; 23 genes) and E (Amino Acid metabolism and transport; 23 genes) (Table A8b).

Most interestingly, *R. tumorigenes* strains carried a gene cluster (*imp*; At1078_04796-At1078_04811) associated with the type VI secretion system (T6SS). This gene cluster was encoded on putative chromid 1 of strains 1078^T^ and 932 (Table A8b). Although they were not identified as species-specific by GET_HOMOLOGUES, *R. tumorigenes* strains carried multiple copies of *vgrG* and single copies of *paar* genes, which are also associated with T6SS machinery (data not shown). *R. tumorigenes* strains also carried a gene cluster (At1078_04243-At1078_04247) annotated as being involved in the synthesis of pseudaminic acid.

On putative chromid 3 of strain 1078^T^, or on putative chromid 2 of strain 932, a putative gene encoding polygalacturonase (glycoside hydrolase family 28) was identified (Table A8b). Polygalacturonase protein sequence of *R. tumorigenes* 1078^T^ shared only 20.7% amino acid identity (69% of query coverage) with polygalacturonase protein of *A. ampelinum* S4^T^ (Avi_1489), which was previously described (Herlache et al. 1997). On the other hand, orthologous protein sequences showing relatively high amino acid identity (>74%) with the polygalacturonase protein sequence of strain 1078^T^ were identified in various members of *Agrobacterium* clade “rubi”, e.g. in *Agrobacterium vaccinii* (84.12% amino acid identity) (Puławska et al. 2022).

## 4 | DISCUSSION

### 4.1 | Novel insights into the taxonomic diversity of agrobacteria

In this study, we conducted polyphasic characterization of “tumorigenes” clade representatives and described a novel species *R. rhododendri* (see below the protologue). The species *R. rhododendri* comprised tumorigenic strains isolated from aerial tumors on rhododendron (Kuzmanović et al. 2019), but also additional strains originating from blueberry and Himalayan blackberry in Oregon (USA) (Weisberg et al. 2022) whose genome sequences were retrieved from GenBank. *R. rhododendri* represents an additional *Rhizobium* species associated with crown/cane gall disease, which further expands our understanding of taxonomic diversity of agrobacteria. By searching the NCBI GenBank (nr/nt and wgs databases), we could not identify additional strains belonging to “tumorigenes” clade. Nevertheless, we assume that members of this clade are distributed more widely and their genetic diversity still remains to be explored.

Apart from *R. rhizogenes* and the “tumorigenes” clade, agrobacteria also occur within other *Rhizobium* clades. In particular, “tropici-rhizogenes” clade comprises at least two additional species that include tumorigenic strains. The first one corresponds to *Rhizobium* sp. AB2/73 which was isolated from *Lippia canescens* in USA (Anderson and Moore 1979). Recently, Hooykaas and Hooykaas (2021) suggested that this strain belongs to a novel *Rhizobium* species. However, our results suggested that this strain most likely belongs to *Rhizobium dioscoreae*, although further taxonomic analysis would help resolve this issue. The second putative species includes Ti plasmid carrying strains 17-2069-2b and 17-2069-2c isolated from blackberry (Weisberg et al. 2022). The closest relative of this potentially novel species is *Rhizobium hainanense* (Figure A5, Table A4).

### 4.2 | Genome architecture of the “tumorigenes” clade

The large number of extrachromosomal elements in members of the “tumorigenes” clade is not surprising as the genomes of nearly all members of the family *Rhizobiaceae* consist of a multipartite architecture (Geddes et al. 2020), with *R. leguminosarum* Rlv3841 containing six extrachromosomal replicons between 151 and 870 kb (Young et al. 2006). Non-chromosomal replicons vary in size and essentiality. While classification systems exist to classify replicons into distinct classes (i.e., plasmid, megaplasmid, chromid), it has been argued that these groups of replicons belong to a spectrum with blurred boundaries (diCenzo and Finan 2017; Hall et al. 2022). We agree with this perspective; yet, we also consider that classification of replicons into distinct groups can nevertheless be useful, in some circumstances, to quickly convey general properties of a replicon of interest.

The genomes of both *R. tumorigenes* strains are split across six replicons: one chromosome, three putative chromids, and two megaplasmids that includes a Ti plasmid. Phylogenetic analysis indicated that the five extrachromosomal replicons of *R. tumorigenes* 1078^T^ had a corresponding replicon in strain 932, as well as corresponding replicons or regions in *R. rhododendri* rho-6.2^T^. In contrast, the related organism *R. tubonense* CCBAU 85046^T^ appears to have two extrachromosomal replicons based on our RepA analysis (Figure S4); however, neither appeared to be orthologous to any of the extrachromosomal replicons of *R. tumorigenes* 1078^T^. We thus conclude that the five extrachromosomal replicons of *R. tumorigenes* were acquired by an ancestor after the split from *R. tubonense* but prior to the split from *R. rhododendri*.

Of the five extrachromosomal replicons of *R. tumorigenes*, three were classified as putative chromids according to the sequence-based classification scheme of diCenzo and Finan (diCenzo and Finan 2017). Chromids generally display higher conservation of gene content than to megaplasmids (diCenzo and Finan 2017). Indeed, compared to the megaplasmids, the putative chromids of *R. tumorigenes* displayed higher conservation both between *R. tumorigenes* strains and with the corresponding replicons or regions of *R. rhododendri* rho-6.2. In addition, as is common for chromids (diCenzo and Finan 2017), the putative chromids appeared to lack conjugation machinery unlike the megaplasmids. Thus, several lines of evidence are consistent with the three putative chromids of *R. tumorigenes* representing true chromids. However, as the defining feature of chromids is that they are essential for cell viability (diCenzo and Finan 2017; Harrison et al. 2010), experimental follow-up is required to definitely classify these replicons as chromids.

*R. rhododendri* rho-6.2^T^ contains two fewer extrachromosomal replicons than do *R. tumorigenes* strains 1078^T^ and 932. Synteny and phylogenetic analyses indicated that this is due to a co-integration of the megaplasmid and two putative chromids in the *R. rhododendri* lineage following divergence from *R. tumorigenes*. Interestingly, two of the three copies of the *repC* gene, encoding plasmid replication proteins, are truncated. The loss of the extra *repC* copies may have helped to stabilize the cointegrant. Although the cointegrant did not fully meet our definition of a chromid (while it exhibited a chromid-like DRA distance from the chromosome of 0.29, the GC content difference compared to the chromosome was >1%), we classified this replicon as a putative chromid as parts of the cointegrant are derived from chromid-like replicons. Although not feasible to test experimentally, it would be interesting to observe whether the megaplasmid portion of the cointegrant evolves chromid-like properties over time.

### 4.3 | Diversification of “tumorigenes” clade

In this study, we identified genes specific for each of the two species the *R. rhododendri* and *R. tumorigenes* by examining the pan-genome of the “tumorigenes” clade. Our objective was to identify potentially adaptive features among species-specific genes, in order to gain a better understanding of the ecological differentiation of these species. We recognize, however, that the availability of genomes for only two *R. tumorigenes* strains is a limitation of this analysis, and that the number of species-specific genes will likely decrease as more genomes become available. Nevertheless, based on the available genomes, the majority of species-specific genes are encoded on putative chromids and megaplasmids, which is in line with previous studies analyzing the *A. tumefaciens* species complex (Lassalle et al. 2017) and *All. vitis* species complex (Kuzmanović et al. 2022b) strains. Although most of the species-specific genes are annotated as encoding hypothetical or poorly described proteins, we could determine putative functions for several genes and gene clusters.

Both *R. rhododendri* and *R. tumorigenes* strains carried a putative gene cluster involved in production of cellulose; however, the former species carried an additional cluster with the same putative function. Both clusters were homologous, but divergent in sequence. *Agrobacterium* and *Rhizobium* spp. were reported to synthetize cellulose (reviewed in (Augimeri et al. 2015; Ross et al. 1991). In *Agrobacterium* spp., production of the exopolysaccharide cellulose is associated with attachment of bacteria to plant surfaces (Matthysse et al. 1981). Although cellulose synthesis was not required for virulence of *Agrobacterium*, cellulose mutants could not firmly attach to host plant, which reduced tumor formation (Matthysse 1983). Similarly, in *R. leguminosarum*, cellulose production is involved in rhizobial attachment to plant roots (Smit et al. 1987). As *R. rhododendri* carries two distinct clusters for cellulose synthesis, if both of them are functional, this species might show enhanced ability to colonize different plant hosts.

Unlike *R. tumorigenes*, *R. rhododendri* carried putative genes associated with the uptake of simple sugars, as well as sugar alcohols, such galactitol and sorbitol. These two sugar alcohols, in addition to mannitol, are widely distributed in angiosperms where they may be involved in response to abiotic and biotic stresses (Moing 2000). The potential ability of *R. rhododendri* to process these compounds could contribute to its environmental adaptation and association with higher plants.

On putative chromid 1, *R. tumorigenes* carried a putative gene cluster associated with T6SS. Homologues genes were not identified in any of the *R. rhododendri* strains. The T6SS is commonly found in plant-associated bacteria and can have diverse genetic architecture (Bernal et al. 2018). A putative gene cluster encoding T6SS in *R. tumorigenes* had identical or similar organization as in other *Rhizobiaceae* strains (Wu et al. 2021). In *A. fabrum*, T6SS is involved in interbacterial competition (Ma et al. 2014; Wu et al. 2019). Accordingly, a T6SS in *R. tumorigenes* might contribute to its competitiveness in plant tissue or rhizosphere.

Furthermore, *R. tumorigenes* strains carried a putative gene cluster implicated in the synthesis of pseudaminic acid. Pseudaminic acid is a microbially produced sialic acid-like sugar involved in glycosylation of flagellin, which plays an essential role in flagella assembly of human pathogenic bacteria such as *Campylobacter jejuni* and *Helicobacter pylori* (reviewed in (Salah Ud-Din and Roujeinikova 2018). In *Sinorhizobium fredii*, pseudaminic acid is a component of the capsular polysaccharide (K antigen) associated with nodulation efficiency on some hosts (Le Quéré et al. 2006; Margaret et al. 2012). Therefore, it is tempting to speculate that the synthesis of pseudaminic acid might be involved in tumorigenesis of *R. tumorigenesis* and plant host invasion.

*R. tumorigenes* strains carried putative chromid-borne gene coding for polygalacturonase, one of the most important enzymes associated with cell wall degradation. It has been reported that *All. vitis* species complex strains are able to produce this enzyme, which plays a role in grapevine root decay (McGuire et al. 1991; Rodriguez-Palenzuela et al. 1991). Different rhizobia (*R. leguminosarum* and *Sinorhizobium meliloti*) were also reported to produce polygalacturonase, for which it was postulated to be involved in the root invasion process (Jimenéz-Zurdo et al. 1996). Accordingly, this putative feature in *R. tumorigenes* could also have a role in degradation of the pectin network that comprises plant cell walls and colonization of particular plant hosts.

### 4.4 | Differentiation of novel *Rhizobiaceae* genera

Based on genus demarcation thresholds defined in our previous study (Kuzmanović et al. 2022a), the core *Rhizobium* superclade should be split in at least three genera. Besides the clade “leguminosarum-etli” (*Rhizobium sensu stricto*), which includes the type species of the genus *Rhizobium* (*R. leguminosarum*), clades “tropici-rhizogenes”, and “tumorigenes” represent candidates for new *Rhizobiaceae* genera. In our opinion, such a division of *Rhizobium* species would require additional genomic or phenotypic evidences, thus revealing factors relevant for biological and ecological diversification of these clades. However, this taxonomic revision was not an objective of this work, and we followed the taxonomic scheme preserving the current structure of the genus *Rhizobium*.

The taxonomic position of *R. tubonense* was not completely clear. This species has relatively high proteome relatedness with both “tumorigenes” and “tropici-rhizogenes” clade representatives. For instance, cpAAI comparisons based on 169 marker proteins yielded values slightly above the threshold for genus demarcation (~86%) in both cases (Kuzmanović et al. 2022a). In core-proteome and pan-genome phylogenetic trees, *R. tubonense* was located on a distant branch, although the “tumorigenes” clade was its closest relative. Taken together, *R. tubonense* might represent an additional candidate for a separate *Rhizobium* genus. Nonetheless, this requires further study, including additional phylogenetic lineages more closely related to *R. tubonense*, which are expected to be discovered in the future.

## 5 | CONCLUSIONS

This study revealed additional genomic and taxonomic diversity of tumorigenic agrobacteria. OGRIs and phylogenomic analyses clearly showed that tumorigenic strains isolated from rhododendron represent a novel species of the genus *Rhizobium* for which the name *Rhizobium rhododendri* sp. nov. is proposed. By searching GenBank, additional *R. rhododendri* strains isolated from blueberry and Himalayan blackberry in USA were identified. Both species of the “tumorigenes” clade (*R. rhododendri* and *R. tumorigenes*), contain multipartite genomes, including a chromosome, putative chromids, and megaplasmids. Interestingly, these two species showed distinct genome architecture. Our investigation indicated that the large putative chromid of *R. rhododendri* is a cointegrant of a *R. tumorigenes*-like ancestral megaplasmid and two putative chromids. Moreover, evidence of inter-replicon DNA exchange between putative chromids of one *R. tumorigenes* lineage was detected. Furthermore, we examined the pan-genome of members of the “tumorigenes” clade and identified genes specific to each of the species *R. rhododendri* and *R. tumorigenes*. For some of the genes and gene clusters, it was possible to determine the putative function and possible role in the ecological adaptation of the studied bacterial species. The predicted functions are found to be primarily associated with plant-bacterial interactions, bacterial competitiveness in plant tissue or rhizosphere, and uptake of specific nutrient sources.

### Description of *Rhizobium rhododendri* sp. nov

*Rhizobium rhododendri* (rho.do.den’dri. N.L. gen. n. *rhododendri*, of *Rhododendron*, the plant genus from which the type strain was isolated).

Bacterial cells are Gram-negative, motile and non-spore forming. They are aerobic, and oxidase and catalase positive. Bacteria grow well on YMA, TY, PDA-CaCO_3_, and R2A media, whereas weak growth was observed on King’s medium B. Colonies on YMA medium had a diameter of 1-2 mm after 72 h of growth at 28°C. They were white to cream colored, circular, convex and glistening. Growth was observed at a temperature range between 5 and 30°C. Nitrate reduction, indole production, and glucose fermentation are negative. Arginine dihydrolase and gelatin hydrolysis tests are negative. Esculin hydrolysis and b-galactosidase tests are positive. D-glucose, D-mannose, and D-mannitol are assimilated. A weak assimilation was observed for L-arabinose and D-maltose. Potassium gluconate, caprate, adipate, malate, trisodium citrate, and phenylacetate are not assimilated. Strain forms clear zones on PDA-CaCO_3_, but do not produce 3-ketolactose from lactose. The major fatty acids (>5%) are C_18:1_ (~6%). *ω*7c (~50%), C_19:0_ cyclo *ω*7c (~19%), C_16:0_ (~7%), and C_18:1_ *ω*7c 11-methyl (~6%).

*R. rhododendri* strains rho-6.2^T^, rho-1.1 and rho-13.1 caused tumors on inoculated rhododendron, sunflower and tomato plants, and were proven to carry a Ti plasmid ((Kuzmanović et al. 2019), this study).

The genome size of the type strain (rho-6.2^T^) is 5.96 Mb. The genome is composed of a circular chromosome (3.71 Mb) and 3 extrachromosomal replicons that are 1.53 Mb, 382 kb, and 337 kb in size. The GC content of the total genomic DNA is 59.98%.

*R. rhododendri* can be distinguished from other *Rhizobium* spp. based on OGRIs (e.g. ANI and dDDH) calculations.

The type strain, rho-6.2^T^ (= DSM 110655^T^ = CFBP 9067^T^) was isolated from an aerial tumor on *Rhododendron* sp. in Germany in 2017. The DDBJ/ENA/GenBank accession numbers for the genome sequence are XX000000 to XX000000 (NCBI submission is undergoing processing and accession numbers will be added when available).

## DATA AVAILABILITY STATEMENT

The whole-genome sequences have been deposited at DDBJ/ENA/GenBank under the accessions XXX (NCBI submission is undergoing processing and accession numbers will be added when available), within the BioProject PRJNA910953.

The raw sequencing reads were deposited in the Sequence Read Archive (SRA) under the same BioProject PRJNA910953.

Other relevant data, including .fasta and .gbk files used for core-genome and pan-genome analyses are available through Figshare (https://figshare.com/) at https://doi.org/10.6084/m9.figshare.21785609, https://doi.org/10.6084/m9.figshare.21785456, https://doi.org/10.6084/m9.figshare.21785570, https://doi.org/10.6084/m9.figshare.21785573, and https://doi.org/10.6084/m9.figshare.21785600. (DOIs will be published upon manuscript acceptance).

## AUTHOR CONTRIBUTIONS

**Nemanja Kuzmanović:** Conceptualization (lead); investigation (leading); formal analysis (lead); data curation (lead); writing – original draft (lead); writing – review and editing (equal); funding acquisition (lead), visualization (equal). **George C. diCenzo:** Conceptualization (equal); formal analysis (equal); data curation (equal); writing – original draft (equal); writing – review and editing (equal); visualization (equal). **Boyke Bunk:** formal analysis (supporting); data curation (supporting); writing – review and editing (equal). **Cathrin Spröer:** investigation (supporting). **Anja Frühling:** investigation (supporting). **Meina Neumann-Schaal:** investigation (equal); writing – review and editing (equal); **Jörg Overmann:** Resources (supporting); writing – review and editing (equal), **Kornelia Smalla:** Conceptualization (supporting); resources (equal); supervision (supporting); writing – review and editing (equal); funding acquisition (suporting).

## ACKNOWLEDGEMENTS

The authors would like to thank Prof. Aharon Oren (The Hebrew University of Jerusalem, Israel) for helpful advice on nomenclatural aspects. This research was enabled, in part, through computational resources provided by BMBF-funded de.NBI Cloud within the German Network for Bioinformatics Infrastructure (de.NBI) (031A537B, 031A533A, 031A538A, 031A533B, 031A535A, 031A537C, 031A534A, 031A532B). We thank Simone Severitt, Jolanthe Swiderski, Nicole Heyer, Anika Wasner and Gesa Martens for excellent technical assistance.

## ETHICS STATEMENT

None required.

## ORCID

Nemanja Kuzmanović 0000-0002-3635-6813

George diCenzo 0000-0003-3889-6570

Boyke Bunk 0000-0002-8420-8161

Meina Neumann-Schaal 0000-0002-1641-019X

Jörg Overmann 0000-0003-3909-7201

Kornelia Smalla 0000-0001-7653-5560

