## Appendix_Figures A1-A10 for "Genomics of the “tumorigenes” clade of the family *Rhizobiaceae* and description of *Rhizobium rhododendri* sp. nov."

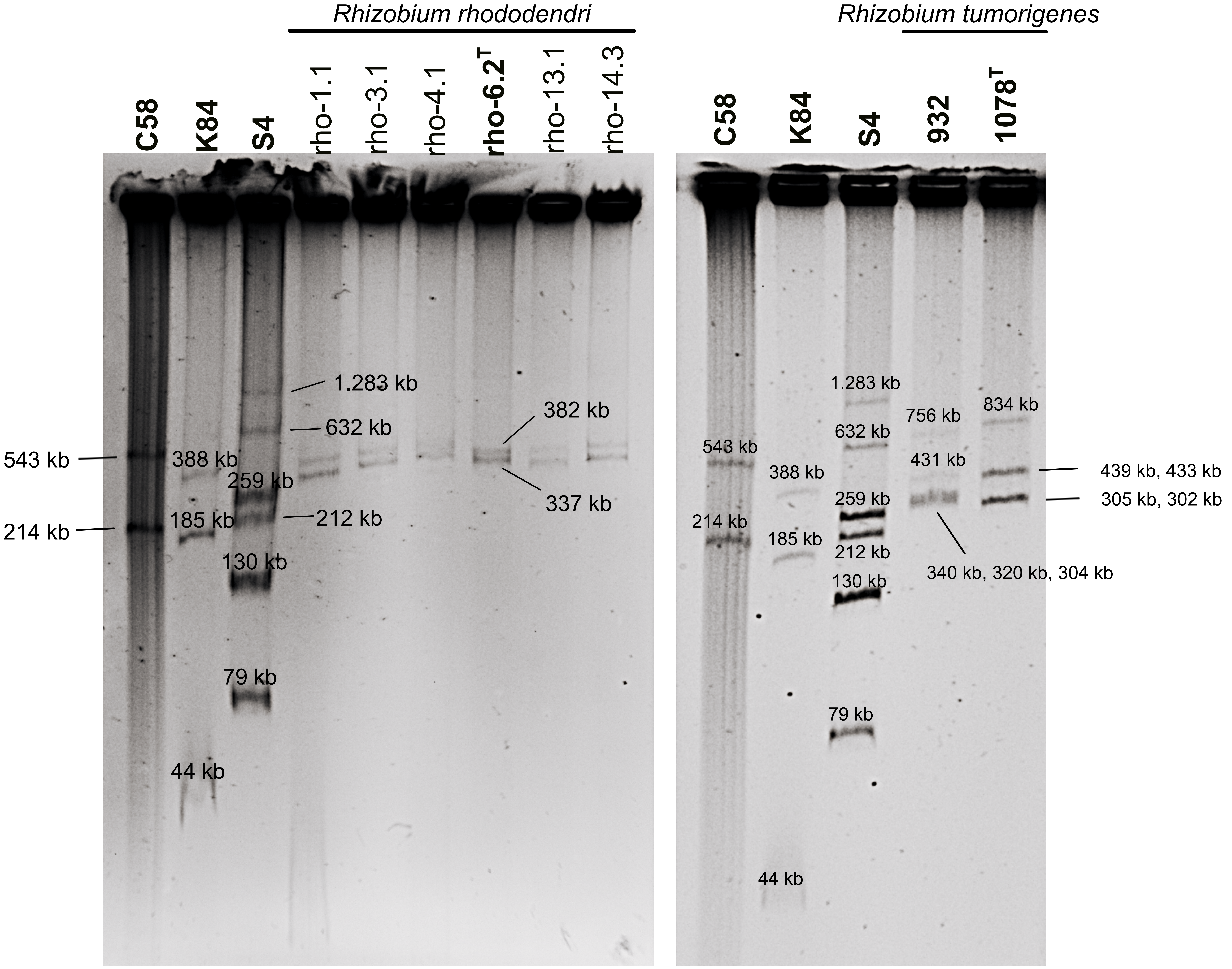


**Figure A1.** Eckhardt-type agarose gel electrophoresis of extrachromosomal replicons of *Rhizobium rhododendri* rho-6.2$^T$, and *Rhizobium tumorigenes* 1078$^T$ and 932. Although finished genomes were not obtained for *R. rhododendri* strains rho-1.1, rho-3.1, rho-4.1, rho-13.1 and rho-14.3 in this study, they were also examined by gel electrophoresis. "*Agrobacterium fabrum*" C58$^T$, *Allorhizobium ampelinum* S4$^T$, and *Rhizobium rhizogenes* K84 carrying replicons of known size were used as markers.

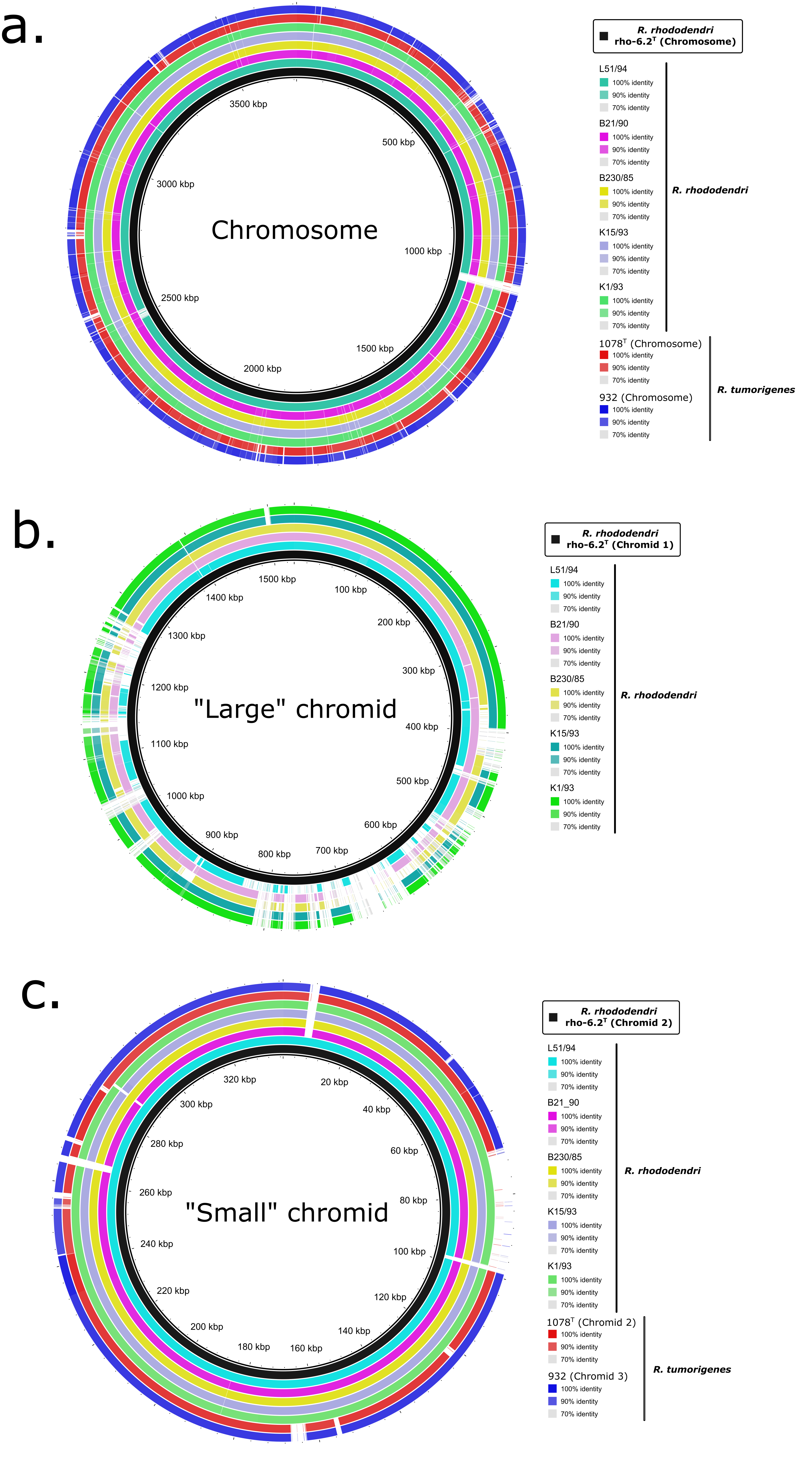


**Figure A2.** Comparison of chromosome (a), chromid 1 (b) and chromid 2 (c) of *Rhizobium rhododendri* rho-6.2$^T$ (reference sequences) with the orthologous replicons of related *R. rhododendri* and *Rhizobium tumorigenes* strains (query sequences). Genetic coordinates of the reference sequences are shown within the thin inner ring. The thick black rings portray corresponding replicons of *Rhizobium rhododendri* rho-6.2$^T$. The colored rings portray query sequences, as indicated in the Figure. Accession numbers of sequences used for comparison are indicated in Table A3. The Figure was generated using BRIG software and edited with Inkscape (see M&M for details).

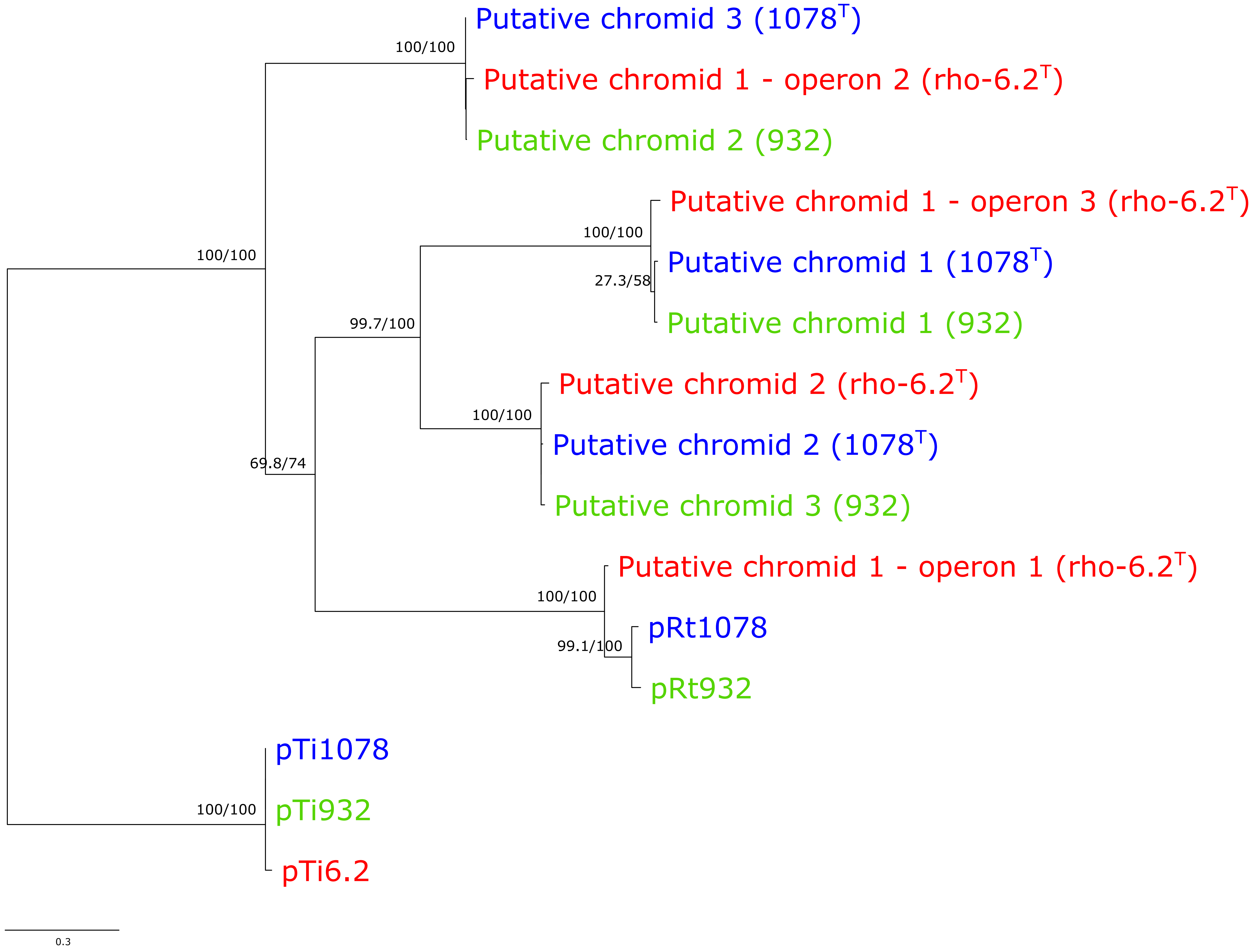


**Figure A3.** Maximum likelihood tree based on the concatenated sequence of RepA and RepB proteins identified in genomes of *Rhizobium rhododendri* rho-6.2$^T$, and *Rhizobium tumorigenes* 1078$^T$ and 932. Corresponding strains are indicated in parentheses or within replicon names. The tree was constructed using model LG+G4. The numbers on the nodes indicate the SH-aLRT support values (first value) and ultra-fast bootstrap values (second value). The tree was midpoint rooted. The scale bar represents the estimated number of amino acid substitutions per site. The trees based on individual RepA and RepB sequences showed identical topology to the tree shown here.

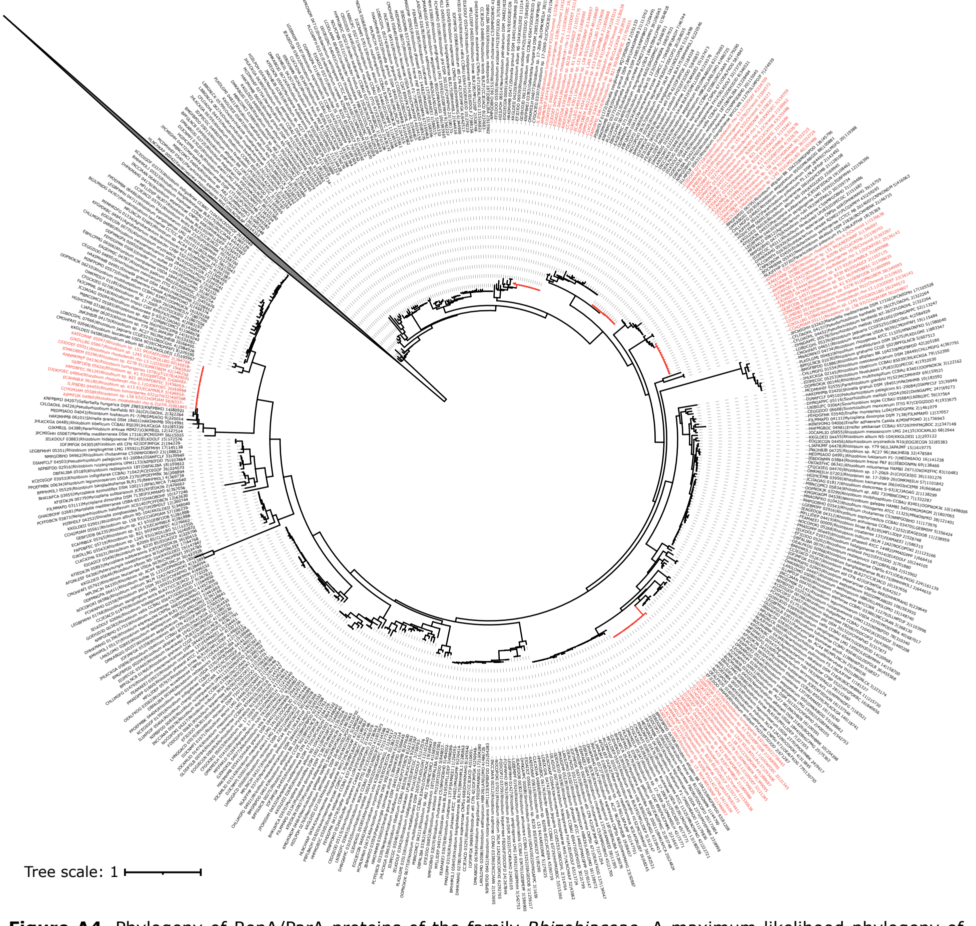


**Figure A4.** Phylogeny of RepA/ParA proteins of the family *Rhizobiaceae*. A maximum likelihood phylogeny of RepA/ParA proteins of 116 *Rhizobiaceae* strains is shown. The extrachromosomal replicons from the "tumorigenes" clade are shown in red. The collapsed clade represents the *Rhizobiaceae* chromosomes. Taxa names are in the format: locus tag | species and strain name | replicon name| replicon size. The scale represents the mean number of amino acid substitutions per site.

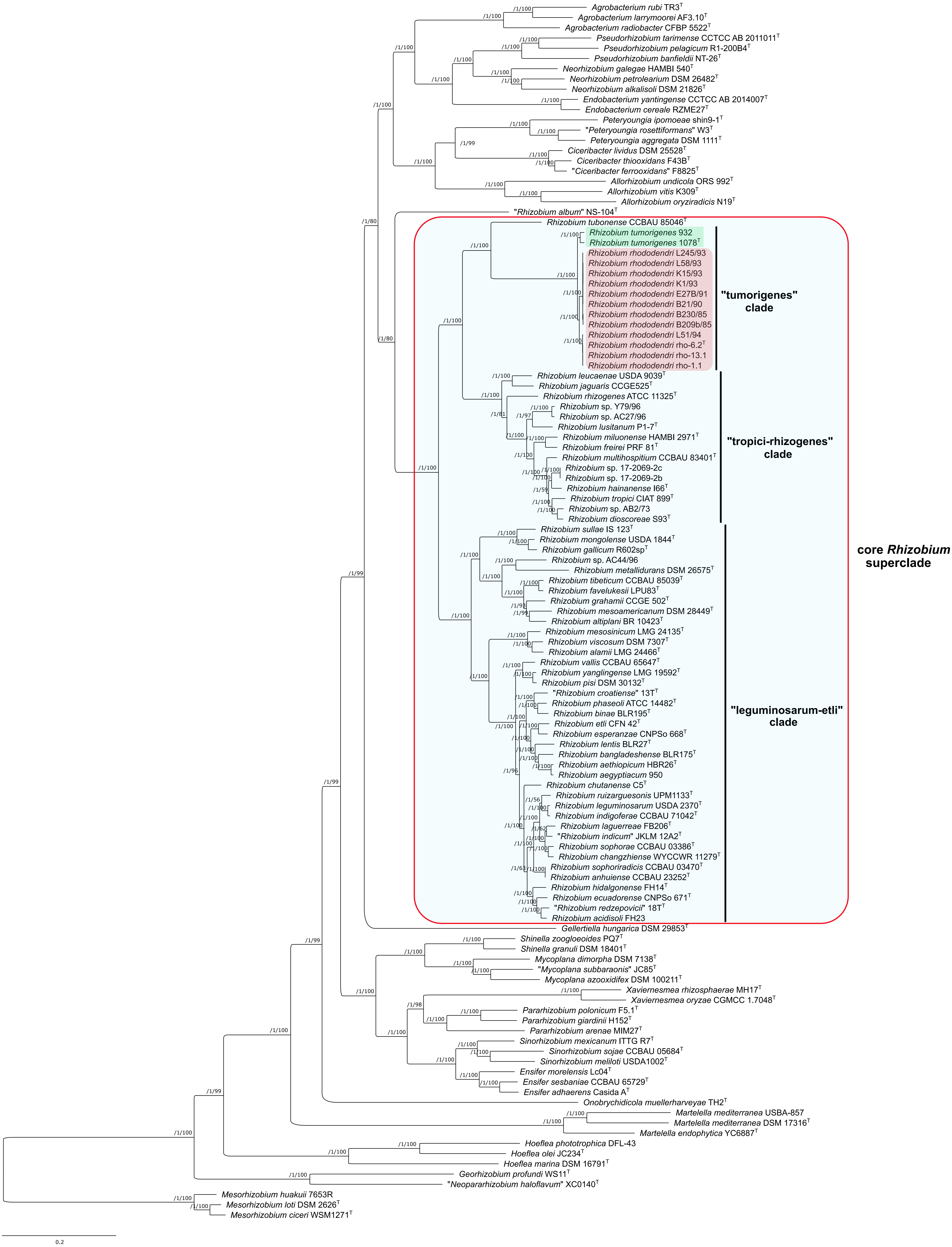


**Figure A5.** Maximum-likelihood core-proteome phylogeny showing the evolutionary relationships between and within the "tumorigenes" clade and other *Rhizobiaceae* clades (uncollapsed). Three *Mesorhizobium* spp. strains were included as the outgroup to root the tree. The phylogeny was estimated from the concatenated alignments of 191 protein sequences selected as top-scoring markers using the GET_PHYLOMARKERS software. The numbers on the nodes indicate the approximate Bayesian posterior probabilities support values (first value) and ultra-fast bootstrap values (second value), as implemented in IQ-TREE. The scale bar represents the number of expected substitutions per site under the best-fitting LG+F+R6 model. The same tree, but with collapsed clades, is presented in Figure 3.

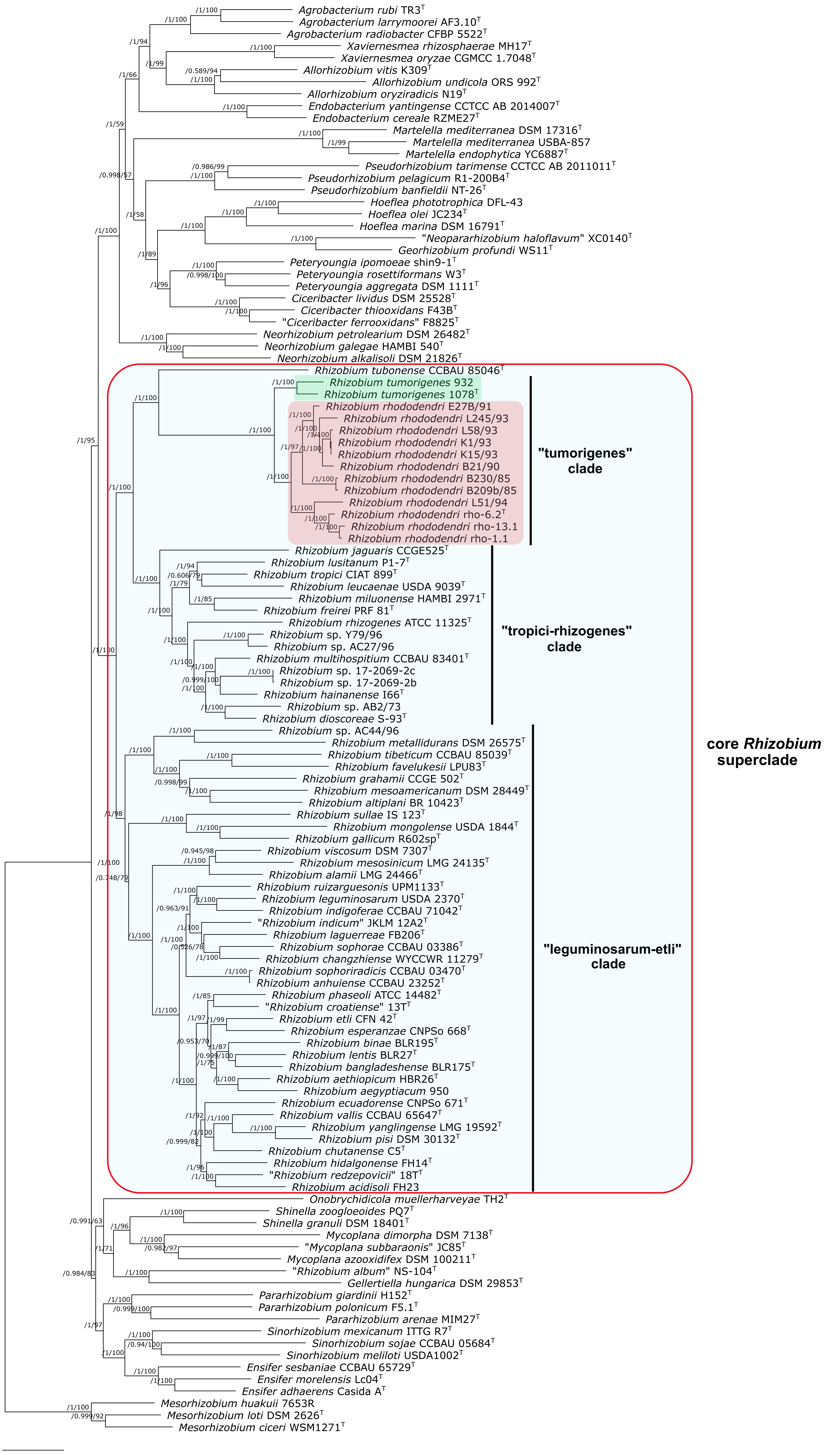


**Figure A6.** Maximum-likelihood pan-genome phylogeny showing the relationships between and within the clade "tumorigenes" and other *Rhizobiaceae* clades (uncollapsed). Three *Mesorhizobium* spp. strains were included as the outgroup to root the tree. The tree was estimated with IQ-TREE from the consensus (COGtriangles and OMCL clusters) gene presence/absence matrix containing 71,538 clusters obtained using GET_HOMOLOGUES software. The numbers on the nodes indicate the approximate Bayesian posterior probabilities support values (first value) and ultra-fast bootstrap values (second value), as implemented in IQ-TREE. The scale bar represents the number of expected substitutions per site under the best-fitting GTR2+FO+R5 model. The same tree, but without collapsing clades, is presented in the Figure 4.

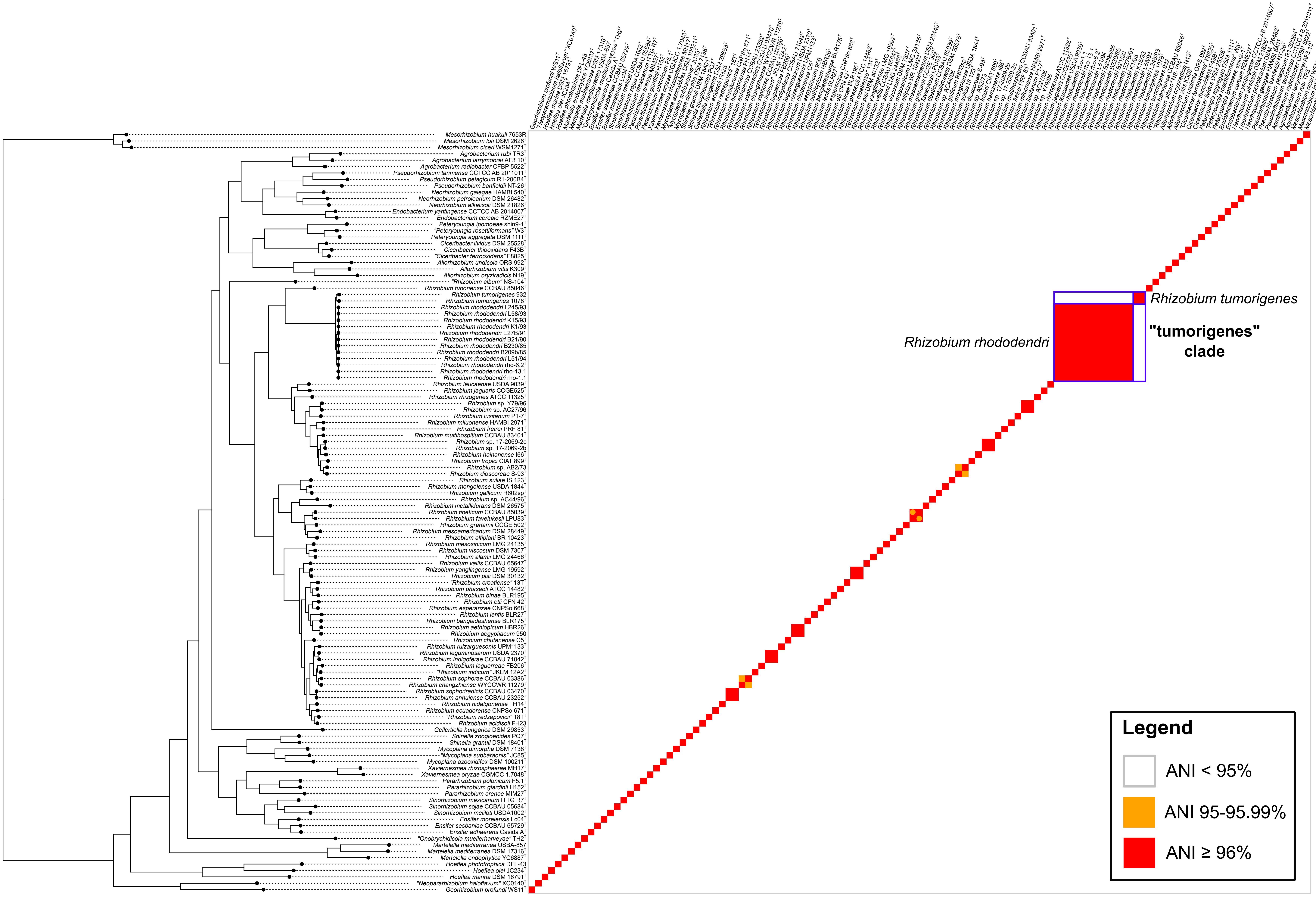


**Figure A7.** Clustered heatmap of average nucleotide identity (ANI) values between the members of the clade “tumorigenes” and other *Rhizobiaceae* clades. Three *Mesorhizobium* spp. strains were included as the outgroup. The ANI calculations were performed using ANIb, OrthoANIu and FastANI methods. The only incongruence between these three methods was observed for strains *Rhizobium favelukesii* LPU83$^{T}$ and *Rhizobium tibeticum* CCBAU 85039$^{T}$, for which the ANIb and orthoANIu values were ≥96%, while the fastANI value was at the threshold for species delineation (95-95.99%), as indicated by orange circles on the heatmap. ANI values were clustered using the core-proteome phylogeny of Figures 3 and A5. The “tumorigenes” clade and its species are indicated with blue boxes.

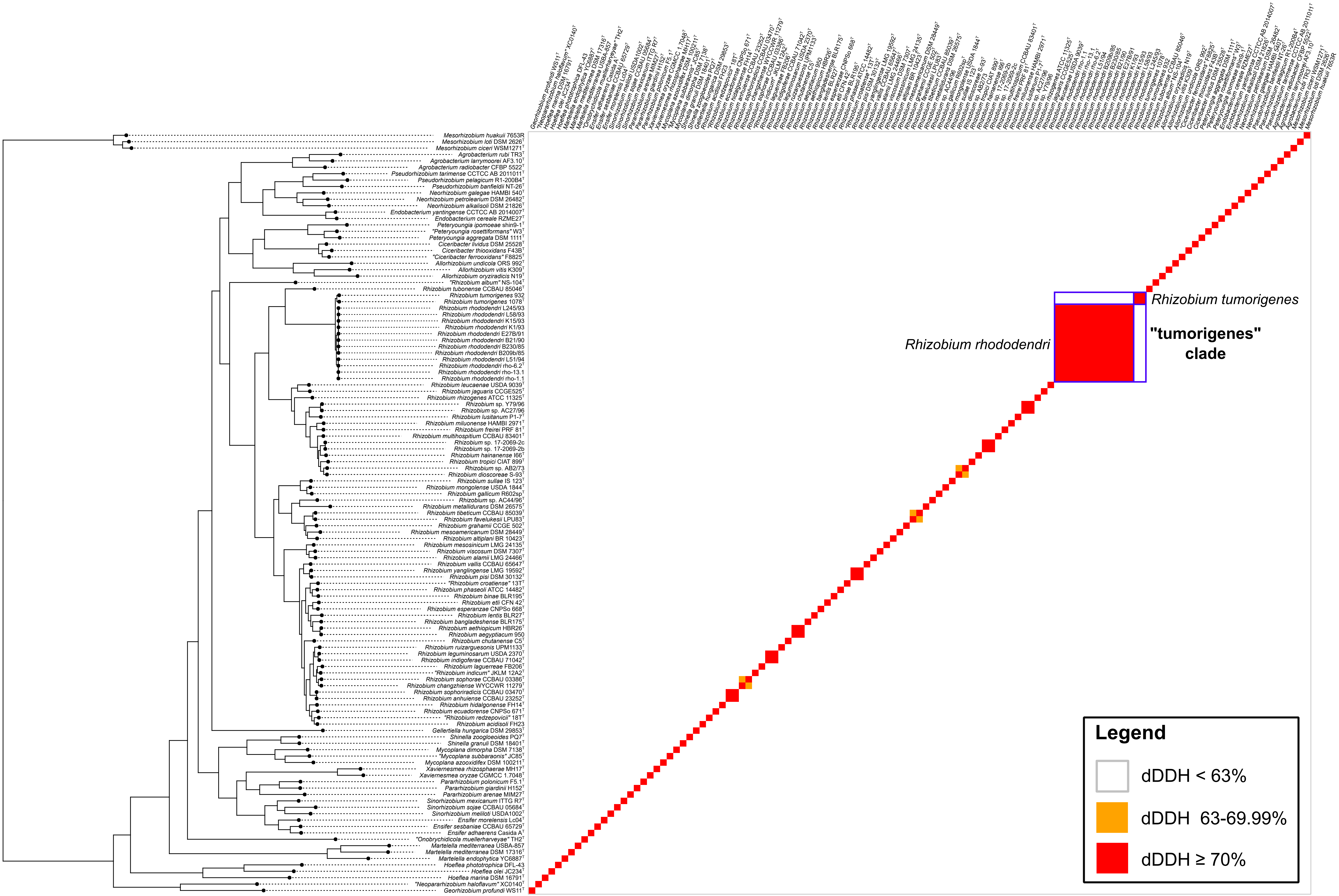


**Figure A8.** Clustered heatmap of digital DNA-DNA hybridization (dDDH) values between the members of the clade “tumorigenes” and other *Rhizobiaceae* clades. Three *Mesorhizobium* spp. strains were included as the outgroup. dDDH values were clustered using the core-proteome phylogeny of Figures 3 and A5. The “tumorigenes” clade and its species are indicated with blue boxes.

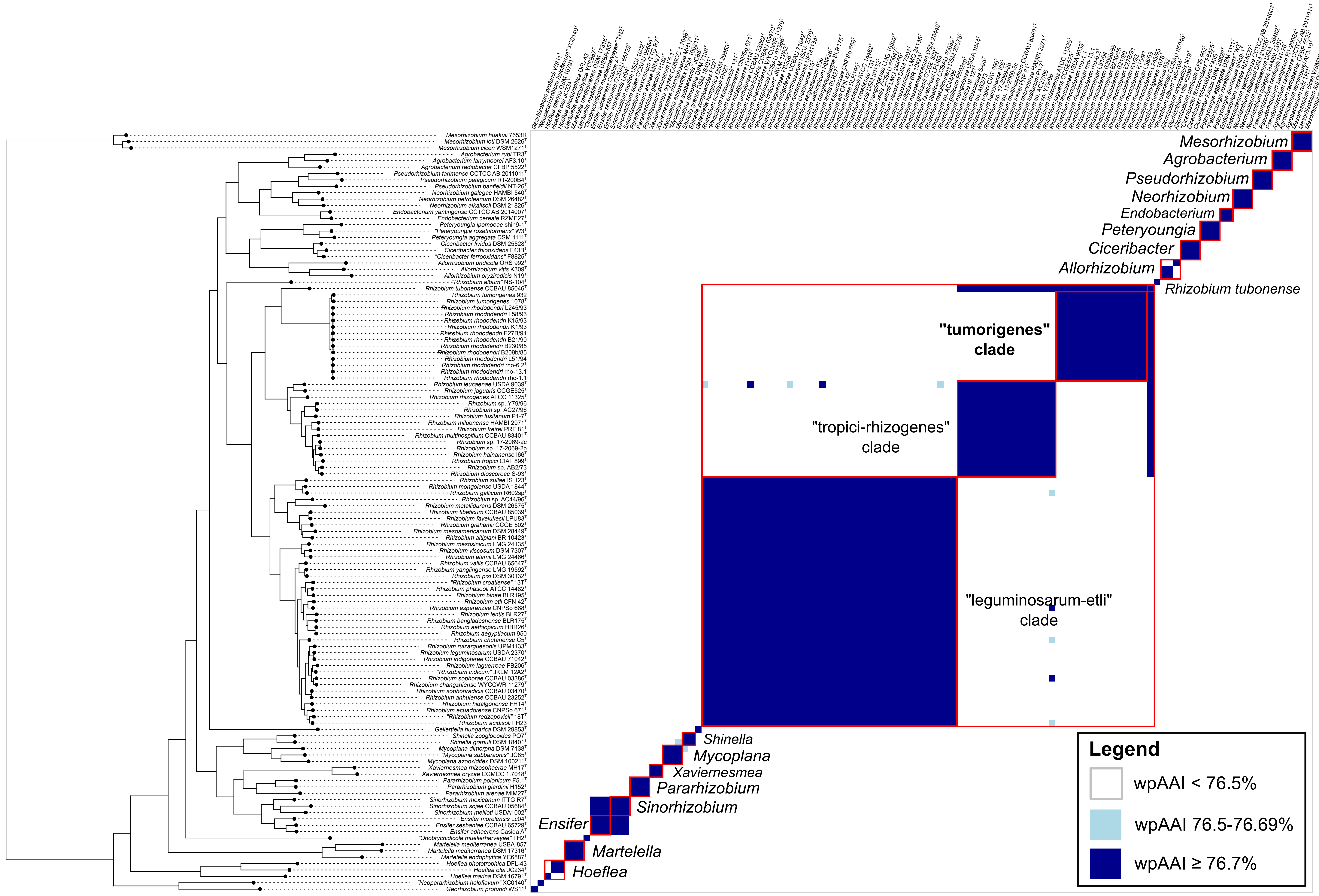


**Figure A9.** Clustered heatmap of whole-proteome average amino-acid identity (wpAAI) values between the members of the clade "tumorigenes" and other *Rhizobiaceae* clades. Three *Mesorhizobium* spp. strains were included as the outgroup. wpAAI values were clustered using the core-proteome phylogeny of Figures 3 and A5. The "tumorigenes" clade and other *Rhizobiaceae* clades are indicated with red boxes.

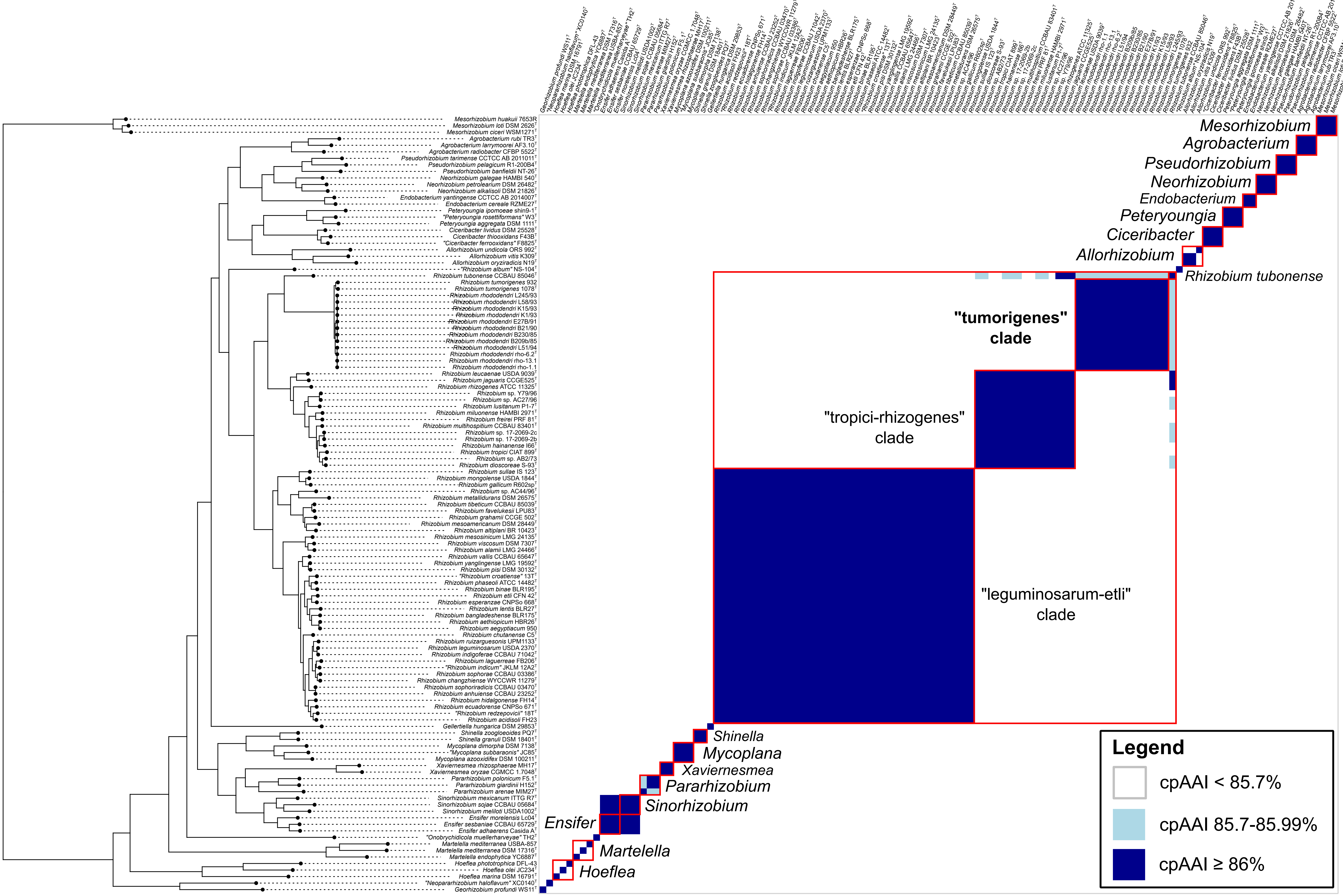


**Figure A10.** Clustered heatmap of core-proteome average amino-acid identity (cpAAI) values between the members of the clade "tumorigenes" and other *Rhizobiaceae* clades. Three *Mesorhizobium* spp. strains were included as the outgroup. cpAAI values were computed from 191 marker proteins selected by GET_PHYLOMARKERS software from the dataset used in this study. These 191 marker proteins were employed to infer core-proteome phylogeny shown in Figures 3 and A5. The core-proteome phylogeny was used to cluster cpAAI values. The "tumorigenes" clade and other *Rhizobiaceae* clades are indicated with red boxes.
