## Supplementary material for "Genomics of the “tumorigenes” clade of the family *Rhizobiaceae* and description of *Rhizobium rhododendri* sp. nov.": Table A5

**Table A5.** Phenotypic characteristics of *Rhizobium rhododendri* rho-6.2^T^ and *Rhizobium tumorigenes* 1078^T^

| **Test** | ***Rhizobium rhododendri* rho-6.2^T^** | ***Rhizobium tumorigenes* 1078^T^** |
| --- | --- | --- |
| Motility | + | + |
| KOH test | + | + |
| Aminopeptidase test | + | + |
| Catalase test | + | + |
| Oxidase test | + | + |
| Growth temperature range (°C) | 5-30 | 5-30 |
| Nitrate reduction^1^ | - | - |
| Indole production^1^ | - | - |
| Glucose fermentation^1^ | - | - |
| Arginine dihydrolase production^1^ | - | - |
| Urease production^1^ | w | + |
| Aesculin hydrolysis^1^ | + | + |
| Gelatin hydrolysis^1^ | - | - |
| β-galactosidase test^1^ | + | + |
| D-glucose assimilation^1^ | + | + |
| L-arabinose assimilation^1^ | w | + |
| D-mannose assimilation^1^ | + | + |
| D-mannitol assimilation^1^ | + | + |
| N-acetyl-glucosamine assimilation^1^ | w | w |
| D-maltose assimilation^1^ | w | w |
| Potassium gluconate assimilation^1^ | - | w |
| Capric acid assimilation^1^ | - | - |
| Adipic acid assimilation^1^ | - | - |
| Malic acid assimilation^1^ | - | w |
| Trisodium citrate assimilation^1^ | - | - |
| Phenylacetic acid assimilation^1^ | - | - |

+, positive; -, negative; w, weak reaction.

^1^ Results obtained with API 20NE system.
