## Supplementary material for "Genomics of the “tumorigenes” clade of the family *Rhizobiaceae* and description of *Rhizobium rhododendri* sp. nov.": Table A6

**Table A6.** Cellular fatty acid composition of *Rhizobium rhododendri* rho-6.2^T^ and *Rhizobium tumorigenes* 1078^T^

| **Fatty Acid^1^** | ***Rhizobium rhododendri* rho-6.2^T^** | ***Rhizobium tumorigenes* 1078^T^** |
| --- | --- | --- |
| C_18:1_ *ω*7c | 49.8 | 51.5 |
| C_19:0_ cyclo *ω*7c | 18.9 | 22.4 |
| C_16:0_ | 6.9 | 5.9 |
| C_18:1_ *ω*7c 11-methyl | 6.3 | 2.4 |
| C_14:0_ 3OH | 4.6 | 4.7 |
| C_16:0_ 3OH | 2.6 | 2.3 |
| C_18:0_ 3OH | 1.9 | 1.6 |
| C_18:1_ *ω*16c | 1.6 | 2.4 |
| C_18:0_ | 1.5 | 1.2 |

^1^ Fatty acids present in amounts lower than 1% and those not confirmed with GC-MS are not shown.
